# Chromatin and transcriptomic profiling uncover dysregulation of the Tip60 HAT/HDAC2 epigenomic landscape in the neurodegenerative brain

**DOI:** 10.1101/2021.02.27.433179

**Authors:** Mariah Beaver, Bhanu Chandra Karisetty, Haolin Zhang, Akanksha Bhatnagar, Ellen Armour, Visha Parmar, Reshma Brown, Merry Xiang, Felice Elefant

**Affiliations:** Department of Biology, Drexel University, Philadelphia, PA 19104, USA

## Abstract

Disruption of histone acetylation mediated gene control is a critical step in Alzheimer’s Disease (AD), yet chromatin analysis of antagonistic histone acetyltransferases (HATs) and histone deacetylases (HDACs) causing these alterations remains uncharacterized. We report the first Tip60 HAT versus HDAC2 chromatin and transcriptional profiling study in *Drosophila* brains that model early human AD. We find Tip60 and HDAC2 predominantly recruited to identical neuronal genes. Moreover, AD brains exhibit robust genome-wide early alterations that include enhanced HDAC2 and reduced Tip60 binding and transcriptional dysregulation. Orthologous human genes to co-Tip60/HDAC2 *Drosophila* neural targets exhibit conserved disruption patterns in AD patient hippocampi. Notably, we discovered distinct transcription factor (TF) binding sites within Tip60/HDAC2 co-peaks in neuronal genes, implicating them in co-enzyme recruitment. Increased Tip60 protects against transcriptional dysregulation and enhanced HDAC2 enrichment genome-wide. We advocate Tip60 HAT/HDAC2 mediated epigenetic neuronal gene disruption as a genome-wide initial causal event in AD.

## INTRODUCTION

Alzheimer’s Disease (AD) is a chronic neurodegenerative disorder affecting the elderly and is the most common cause of dementia. The disease is hallmarked by amyloid-β (Aβ) plaque accumulation, Tau mediated neurofibrillary tangles, and neuronal cell death in the brain that is accompanied by debilitating cognitive deficits in AD patients that worsen as they age. The severity and speed of AD progression are dependent upon complex interactions between genetics, age, and environmental factors (Karch, Cruchaga, & Goate, 2014; Masters et al., 2015; Sanchez-Mut & Graff, 2015), all of which are orchestrated, at least in part, by epigenetic histone acetylation mediated gene control mechanisms. Indeed, decreased chromatin histone acetylation levels have been reported in the brains of animal models and human patients that have multiple types of neurodegenerative diseases that include AD (Berson, Nativio, Berger, & Bonini, 2018; Saha & Pahan, 2006). These alterations have been shown to cause an epigenetic blockade of neuroplasticity gene transcription that contributes to cognitive impairment (Graff et al., 2012; Panikker et al., 2018). More recently, a compelling study using the brains of AD patients reported an age-associated genome-wide reduction of the histone acetylation H4K16 modification that is proposed to contribute to epigenetic gene alteration mediated neurodegeneration (Nativio, Donahue, Berson, Lan, Amlie-Wolf, Tuzer, Toledo, Gosai, Gregory, & Torres, 2018). Despite these informative findings, to date, all AD-associated genome-wide epigenetic studies are limited to examining chromatin histone acetylation patterns and alterations already generated. Thus, little is known about the genome-wide distribution of the antagonizing histone acetyltransferase (HAT) and histone deacetylase (HDAC) enzymes that act to modify the neural epigenome by generating and erasing specific cognition-linked acetylation marks, respectively, and thus serve as the causative agents of memory-impairing histone acetylation alterations in AD.

Appropriate histone acetylation homeostasis in the brain is maintained by HATs and HDACs that in general, activate and repress neural gene expression profiles, respectively. Disruption of this finely tuned HAT and HDAC epigenetic balance causes transcriptional dysregulation that is a key step in AD etiology (Graff et al., 2012; X. Lu, Wang, L., Yu, C., Yu,D., and Yu, G., 2015; Saha & Pahan, 2006; Sanchez-Mut & Graff, 2015). In support of this concept, we and others have reported reduced HAT Tip60 (KAT5) (Panikker et al., 2018) and enhanced HDAC2 (Graff et al., 2012) recruitment to a set of critical neuroplasticity genes in AD animal models and human patients that causes reduced histone acetylation at these gene loci with concomitant transcriptional repression. Nevertheless, whether similar alterations of Tip60 and HDAC2 chromatin distribution with concomitant transcriptional dysregulation are a genome-wide phenomenon that occurs as an early initial event in AD progression remains unknown.

Here we report the first genome-wide study profiling Tip60 HAT versus HDAC2 chromatin distribution and transcriptional dynamics in the brains of amyloid precursor protein (APP) *Drosophila* larvae that effectively model early human AD neurodegeneration both epigenetically and pathologically. We find that Tip60 and HDAC2 predominantly recruited on identical neuronal genes with enrichment peaking across entire gene bodies. Astoundingly, prior to amyloid-β accumulation, AD larval brains exhibit robust genome-wide binding disruptions: enhanced HDAC2 and reduced Tip60 binding with concomitant transcriptional dysregulation. Orthologous human genes to co-Tip60/HDAC2 AD-associated neural targets identified in *Drosophila* exhibit conserved disruption patterns in the human AD hippocampus. Notably, we discovered eight transcription factors (TFs) binding close or within Tip60/HDAC2 co-peaks in neuronal genes, implicating them in co-enzyme recruitment to these loci. Strikingly, increased Tip60 protects against transcriptional dysregulation and enhanced HDAC2 enrichment genome-wide. Based on these results, we advocate that Tip60 HAT/HDAC2 mediated epigenetic transcriptional dysregulation is a genome-wide initial causal event in the AD brain that can be reversed by restoring Tip60/HDAC2 balance.

## RESULTS

### Tip60 protects against early and late transcriptome-wide alterations in the AD-associated neurodegenerative brain

Mild cognitive impairment (MCI) is a debilitating hallmark during early pre-clinical stages of AD, yet the molecular events that trigger these impairments are unclear. We and others have shown that such preclinical AD pathologies in humans are conserved in the well-characterized AD-associated human amyloid precursor protein (APP) *Drosophila* model that inducibly and pan-neuronally express human APP (Fossgreen, Brückner, et al., 1998; Panikker et al., 2018). Third-instar larvae that model early staged APP-induced neurodegeneration show deficits in cognitive ability and synaptic plasticity, axonal transport and outgrowth, and apoptotic neuronal cell death in the brain (Johnson, Sarthi, Pirooznia, Reube, & Elefant, 2013; Panikker et al., 2018; Pirooznia et al., 2012). APP flies also display Aβ plaque accumulation in the aged adult fly eye *via* human conserved endogenous gamma (Fossgreen, Bruckner, et al., 1998) and beta-secretase cleavage pathways (Greeve, 2004). Thus, we first asked whether APP flies also display Aβ plaque formation in the fly brain and whether its accumulation is associated with the early pre-clinical AD defects modeled during larval stages. We focused our studies on the mushroom body (MB) Kenyon cell region as we have shown that Tip60, robustly produced in the MB, is required for MB role in learning and memory and that MB morphology is disrupted in the aged seven-day-old APP fly brain (Xu et al., 2014). Anti-Aβ immunofluorescence studies (Iijima et al., 2008; Iijima et al., 2004) revealed that APP expression in the *Drosophila* brain results in diffuse amyloid deposits that appear in the MB of seven-day-old flies (Fig. 1Aii and 1Bii). These Aβ plaque deposits are unobservable in an earlier AD stage modeled in third-instar larvae (Fig. 1Ai and 1Bi). These results suggest that molecular mechanisms distinct from Aβ plaques trigger early AD pre-clinical impairments.

**Figure 1.**
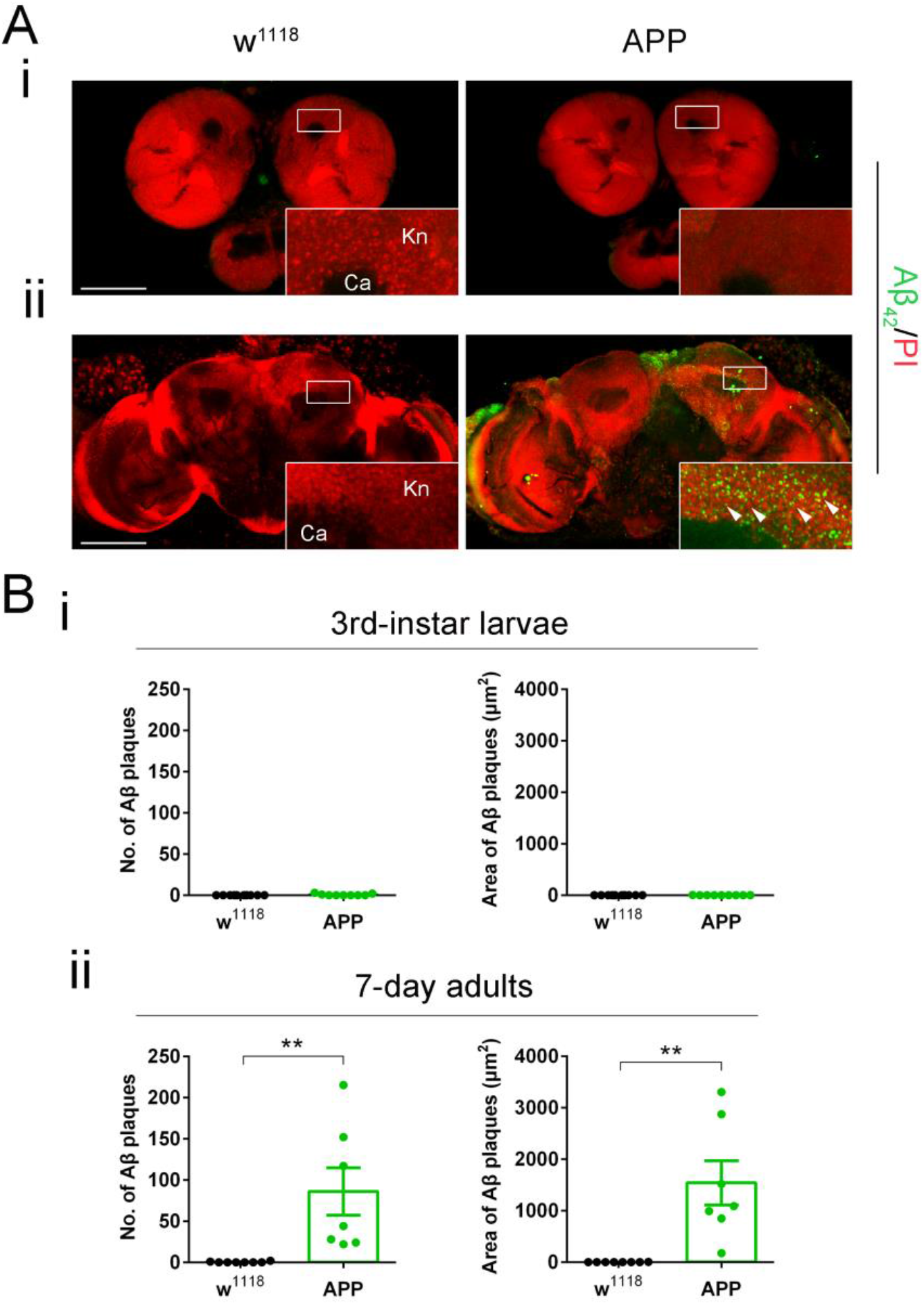
Diffuse amyloid deposits are abundant in the mushroom body (MB) in 7-day APP adults but not in 3rd-instar APP larvae. (A) Representative images. Aβ plaques were stained with anti-Aβ42 antibody (green). Nuclei were stained with PI (red). The Kenyon (Kn) cell region (boxed) was zoomed in to display Kn cells and Aβ plaques. (i) Immunostaining of brains of 3rd-instar larvae shows a negligible Aβ42 signal in APP flies compared to no Aβ42 signal in w1118 flies. (ii) Immunostaining of brains of 7-day adults shows evident Aβ plaques in APP flies compared to w1118 flies. Arrowheads indicate Aβ plaques. No Aβ42 signal was detected in the Calyx (Ca) region. Scale bar represents 100 μm. (B) Aβ plaque was quantified by both number and size. (i) Quantification of Aβ plaque numbers and areas in the 3rd-instar larval brain Kn region. n = 9 ~ 10. (ii) Quantification of Aβ plaque numbers and area in the 7-day adult brain Kn region. n = 8 ~ 9. **p < 0.01; unpaired student’s t-test. All data are shown as mean ± s.e.m.

Gene expression (Grothe et al., 2018; Patel, Dobson, & Newhouse, 2019) and genetic variation (Karch et al., 2014; Kunkle et al., 2019) studies in AD patients and animal models indicate that alteration in gene control contributes to disease pathology. Nevertheless, whether genome-wide gene expression alterations trigger MCI before Aβ plaque formation remains to be further elucidated as gene studies predominantly rely on aged AD brain samples. To address this question, we profiled genome-wide transcriptional changes during early neurodegeneration stages modeled in APP larval brains and later stages modeled in the aged seven-day-old APP fly heads. As we previously identified disruption of Tip60/HDAC2 mediated neuronal gene control as a potential early mechanism underlying neuronal deficits in APP flies (Panikker et al., 2018), we also asked whether increasing Tip60 HAT activity would protect against potential genome-wide early and late-stage gene alterations.

For transcriptome analysis, RNA was isolated from the brains of staged third-instar larvae and from the heads of seven-day-old flies that were w^1118^ control flies or flies expressing either APP or APP;Tip60 under the control of the pan-neuronal elav-GAL4 driver. We used RNA-Seq to quantify gene expression changes. PCA analysis (Supplemental Fig. 1A & 1B) and hierarchically clustered heatmaps (Supplemental Fig. 1C & 1D) show homogeneity within replicates and variability between groups. Importantly, in both early and late developmental stages, the APP;Tip60 transcriptome displays more similarity to the w^1118^ transcriptome than the APP transcriptome (Supplemental Fig. 1C & 1D). Further, tissue enrichment with the human orthologs of the top 2000 APP-induced gene alterations underscores the neural specificity in gene expression defects (Supplemental Fig. 1E & 1F). Reflecting the plaque formation in adult brains, significant alterations in gene expression were identified in the adult APP fly heads (APP vs w^1118^: 1493 up/1641 down). Surprisingly, in the absence of plaque formation in the early APP larval stage brain, we observed even greater changes in gene expression (APP vs w^1118^: 1750 up/2261 down) when compared with adult APP fly heads (Fig. 2A and Supplemental Table 1: S1-1 & S1-3). Consistent with our prior findings demonstrating Tip60 protection against AD defects modeled in APP flies, Tip60 expression led to notable gene expression alterations in both APP larval (APP;Tip60 vs. APP: 311 up/338 down) and adult heads (APP;Tip60 vs. APP: 1023 up/1280 down) (Fig. 2A and Supplemental Table 1: S1-2 & S1-4). We next analyzed the data with the goal of identifying Tip60 rescued genes and associated biological processes specifically reprogrammed by Tip60. To this end, we analyzed the distribution and intersection between down and up-regulated genes between APP vs. w^1118^ and APP;Tip60 vs. APP in larval (Fig. 2B) and adult (Fig. 2C) stages. In the APP larval brain, approximately 11% (458/4011) of gene changes (APP vs w^1118^) are specifically protected against by increased Tip60 and are referred to here as “Tip60 reprogrammed genes”. In APP adult heads, approximately 60% (1898/3134) of APP-induced genes (APP vs w^1118^) were identified as Tip60 reprogrammed genes. Thus, the number of Tip60 rescued genes is significantly greater in the adult stage than in the larval stage. GO analysis revealed that among the top 25 biological processes associated with the Tip60 reprogrammed genes identified in adult flies, axon and dendrite related pathways were enriched (Fig. 3D and Supplemental Table 2: S2-3), while cell-cycle regulation processes and RNA metabolic processes were enriched in the larval stage (Fig. 3A and Supplemental Table 2: S2-2). Lipid metabolic pathways were enriched for Tip60 rescued genes in both adult and larval stages (Fig. 3B & 3C and Supplemental Table 2: S2-4 & S2-1). Our transcriptomic analysis reveals that Tip60 protects against genome-wide gene expression alterations important for neuronal function during early and late-stage AD-associated neurodegeneration with enhanced protection during later stages.

**Figure 2:**
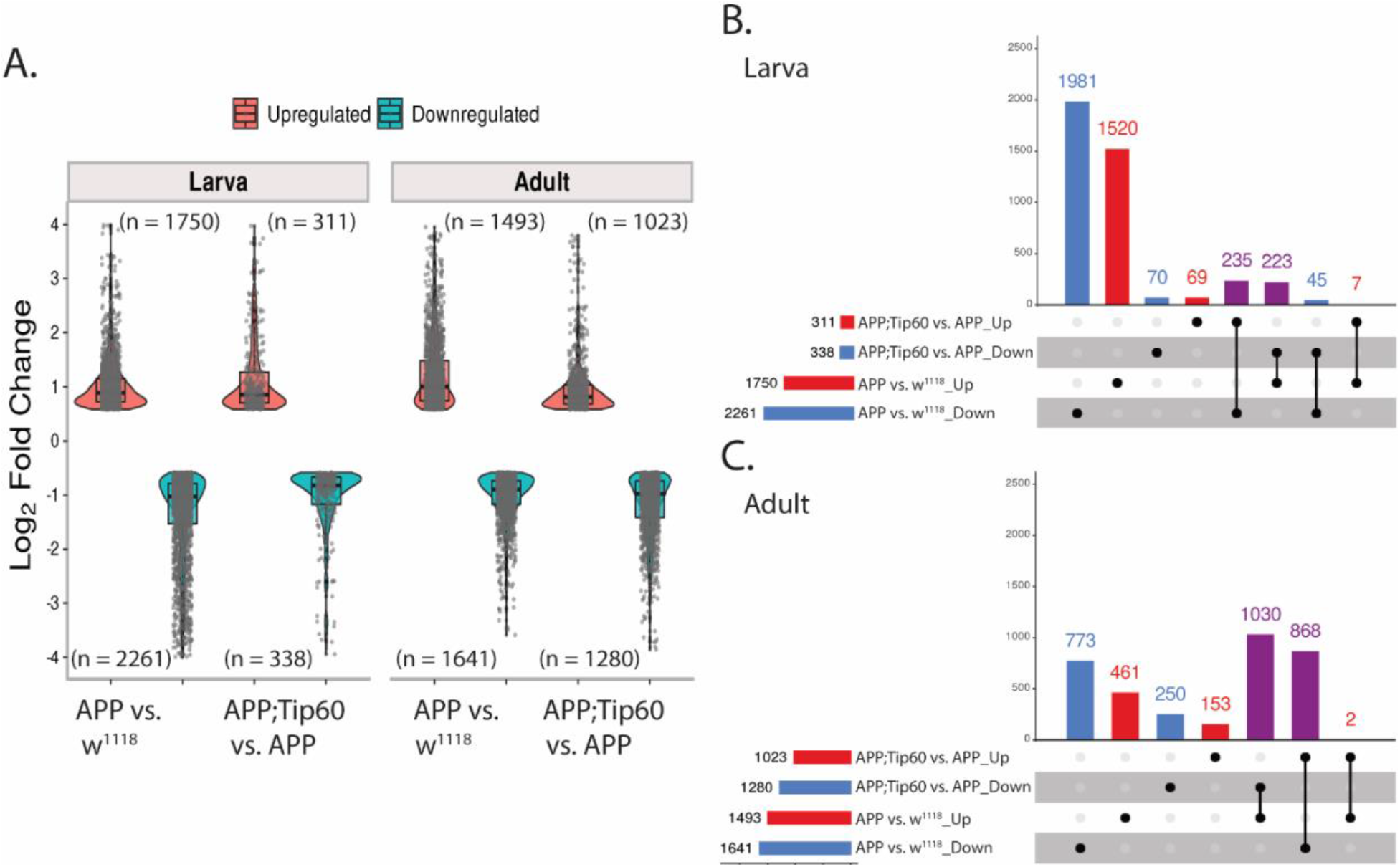
Tip60 protects against early (third instar larval) and late (seven-day-old adult) transcriptomic deregulation in the APP AD associated neurodegenerative brain. (A) Log2 fold changes of differentially expressed genes (padj ≤ 0.05 and log2FoldChange of ≤ −0.583 and ≥ 0.583) determined by RNA-seq in the third instar larval and adult heads in APP vs. w1118 and APP;Tip60 vs. APP. Changes were prominent in both third instar larval and adult APP heads, while Tip60-induced changes initiated in the third instar larval head and were prominent in the adult head: indicating the effect of Tip60 over time. (B & C) The upSet plot represents the distribution and intersection of down and up-regulated genes between APP vs. w1118 and APP;Tip60 vs. APP in third instar larval (B) and adult (C) heads. Rows represent the number of genes in each comparison (APP vs. w1118 and APP;Tip60 vs. APP), and columns represent the number of genes per interaction. The red and blue bars represent the up and down-regulated genes, respectively. The black filled dots indicate the association between rows. The red and blue columns represent genes uniquely up-regulated and down-regulated genes, respectively, in given comparisons, while the purple columns represent Tip60 reprogrammed genes.

**Figure 3:**
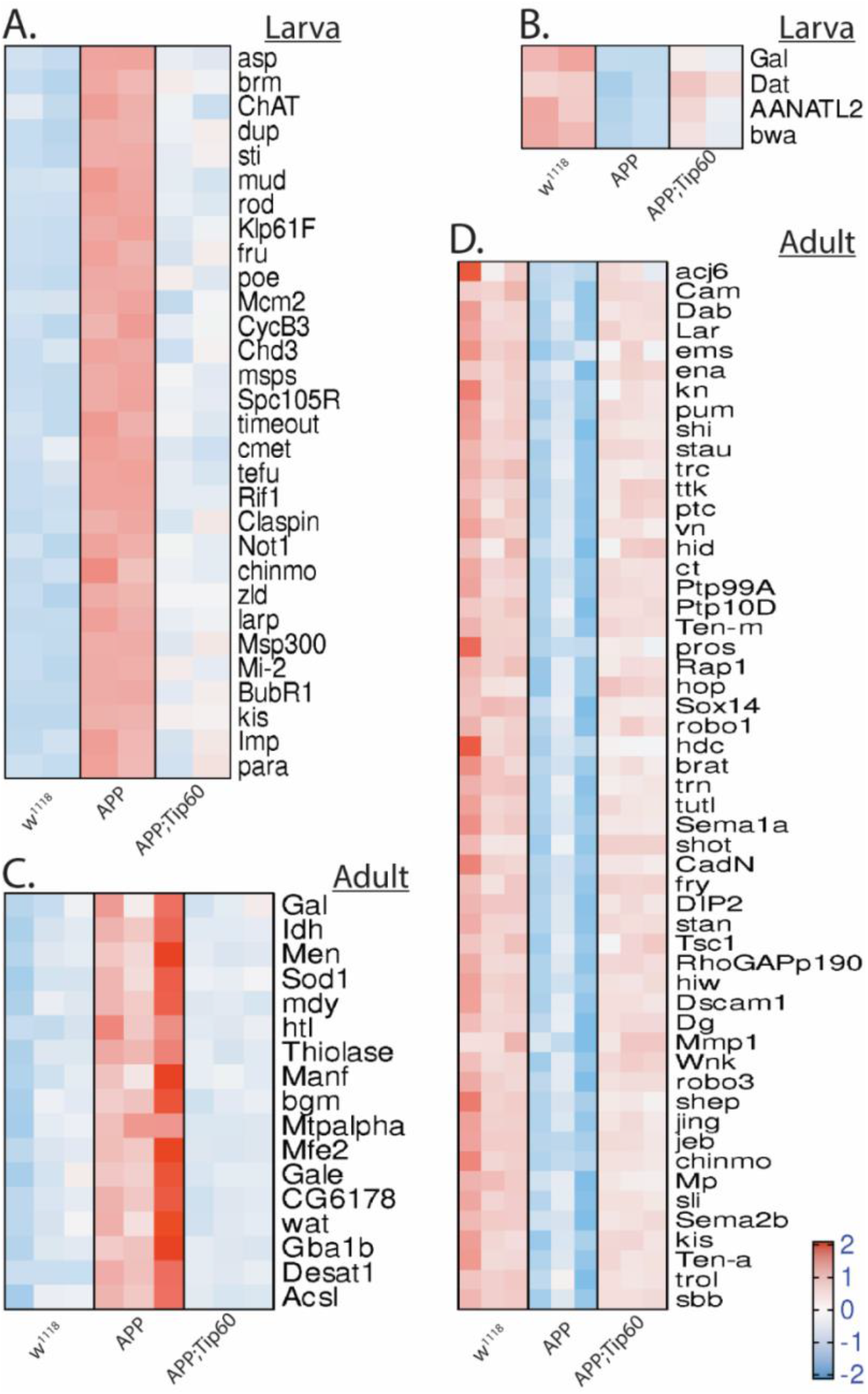
Heatmaps depicting the relative expression pattern of genes misregulated in APP larval and adult heads and are rescued by Tip60. Representation of genes from the most representative biological processes in the top 25 pathways enriched from the rescue gene list. (A) Heatmap of genes representing the cell-cycle regulation processes and RNA metabolic processes in the third instar larval head. Heatmap of genes representing the lipid metabolic pathways in the (B) third instar larval head and (C) the adult head. (D) Heatmap of genes representing the axon and dendrite related pathways in the adult head. Log-transformed gene expression values are displayed as colors ranging from red to blue, as shown in the key. Red represents an increase in gene expression, while blue represents a decrease in expression.

### Increased Tip60 protects against enhanced repressor HDAC2 recruitment along the neuronal gene bodies during early AD neurodegeneration

To elucidate the role of Tip60 and HDAC2 in the early transcriptional dysregulation we observed in the larval brain prior to Aβ plaque formation, we profiled genome-wide enrichment of Tip60 and HDAC2 by ChIP-Seq in larval heads obtained from w^1118^, APP, or APP;Tip60 genotypes (Supplemental Table 3). The peaks identified by ChIP-seq (Supplemental Fig. 2A & 2D) were first annotated to gain insight into their distribution over the genome (Supplemental Fig. 2B & 2E). Interestingly, approximately 60% of the peaks identified for both HDAC2 and Tip60 enrichment were along the gene body (exon and intron regions). A comparative analysis of the genes associated with these peaks among the genotypes (APP, APP;Tip60, w^1118^) revealed ~79% commonality for HDAC2 and ~88% commonality for Tip60 (Supplemental Fig. 2C & 2F). Further, PCA analysis (Supplemental Fig. 3A & 3B) and hierarchically clustered heatmaps (Supplemental Fig. 3C & 3D) shown homogeneity within replicates and variability between groups in both HDAC2 and Tip60. These results suggest that similar genes were regulated in each genotype by Tip60 or HDAC2 and Tip60-induced enrichment in APP;Tip60 was more similar to w^1118^.

We next performed enrichment quantification of the identified Tip60 and HDAC2 ChIP-Seq peaks in w^1118^ control, APP and APP;Tip60 larval heads to determine whether their chromatin binding was altered in APP larval heads (APP vs w^1118^) and whether increased Tip60 could protect against potential binding changes (APP;Tip60 vs. APP). Our findings revealed that in the APP larval heads, there were robust changes in binding enrichment for both HDAC2 (5400 peaks with increased binding and 6571 peaks with decreased binding) and Tip60 (1562 peaks with increased binding and 2023 peaks with decreased binding) (Fig. 4A and Supplemental Table 1: S4-3 & S4-1). Also, tissue enrichment of the top 2000 APP-induced peak enrichment unveils the HDAC2 and Tip60 neural specificity (Supplemental Fig. 3E & 3F). Increased Tip60 levels induced a significant reduction in HDAC2 binding (2718 peaks with increased binding and 8960 peaks with decreased binding) and minimal changes in Tip60 binding (4 peaks with increased binding and 1 peak with decreased binding) (Fig. 4A and Supplemental Table 1: S4-4 & S4-2). This Tip60 mediated trend in reduced HDAC2 binding is evident by the change in the ratio of decreased binding to increased binding (5.5:4.5 in APP vs w1118 to 7.7:2.3 in APP;Tip60 vs. APP): an increase in the number of peaks with decreased binding (6571 in APP vs w^1118^ to 8960 in APP;Tip60 vs. APP) decrease in the number of peaks with increased binding (5400 in APP vs w^1118^ to 2718 in APP;Tip60 vs. APP), and a decrease in the median of log_2_FoldChange with increase in HDAC2 binding.

**Figure 4:**
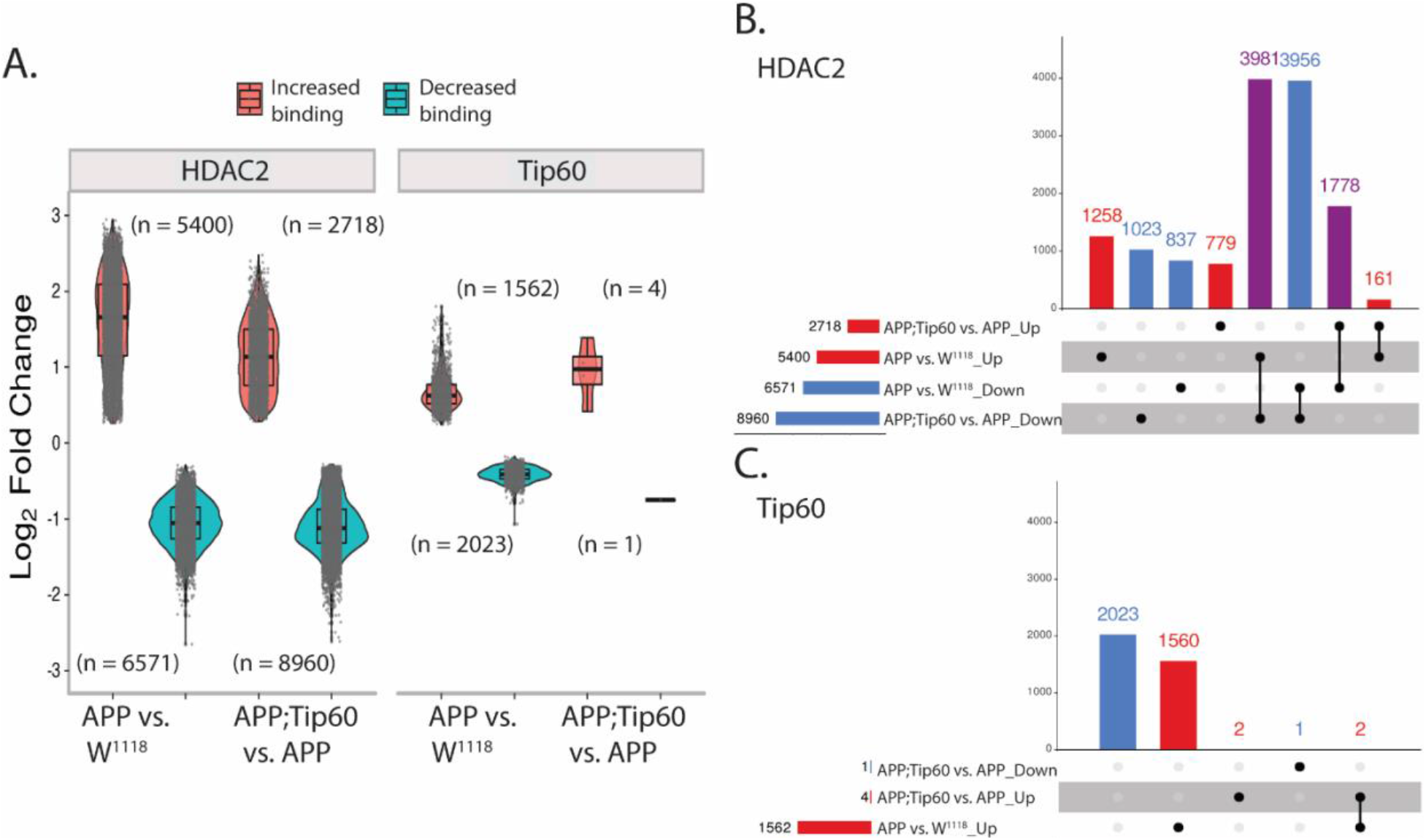
Increased Tip60 protects against enhanced HDAC2 enrichment in APP larval heads. (A) Log2 fold changes of differentially bound peaks (padj ≤ 0.05) of HDAC2 and Tip60 in APP vs. w1118 and APP;Tip60 vs. APP. APP-induced changes (APP vs. w1118) were prominent in both HDAC2 and Tip60 samples, while Tip60-induced changes (APP;Tip60 vs. APP) were prominent only in HDAC2 samples. (B & C) The upSet plot represents the distribution and intersection of differentially bound peaks between APP vs. w1118 and APP;Tip60 vs. APP from HDAC2 (B) and Tip60 (C) samples. Rows represent the number of peaks in each comparison (APP vs. w1118 and APP;Tip60 vs. APP), and columns represent the number of peaks per interaction. The red and blue bars represent the increased and decreased binding of HDAC2 or Tip60, respectively. The black filled dots indicate the association between rows. The red and blue columns represent peaks unique to a given comparison, while the purple columns represent the peaks rescued by Tip60 expression.

We next analyzed the distribution and intersection of altered peaks in larval heads between genotypes APP vs. w^1118^ and APP;Tip60 vs. APP for HDAC2 (Fig. 4B) and Tip60 (Fig. 4C). Remarkably, for HDAC2 binding, approximately 48% (5759/11971) of the total number of peaks altered in APP larval head (APP vs w^1118^) were restored by an increase in Tip60 levels (APP;Tip60 vs. APP). Thus, we refer to these peaks as “Tip60 reprogrammed HDAC2 peaks”. The Tip60 reprogramming effect was primarily observed for HDAC2 binding, visualized with both profile plots (Fig. 5 E i & F i) and heatmaps (Fig. 5 E ii & F ii). As 60% of the identified Tip60 and HDAC2 peaks were enriched along the gene body, we visualized the ChIP-Seq read densities of the significantly altered peak enrichment + /− 0.5 kilobase from the center region of the gene body (Fig. 5 and Supplemental Fig. 4). In APP larval heads, (Fig. 5A-D), the increase in binding of HDAC2 (Fig. 5B) and decrease in binding of Tip60 (Fig. 5C) highly predominates over the decrease in binding of HDAC2 (Fig. 5A) and increase in binding of Tip60 (Fig. 5D). Further, increased Tip60 levels protected against alterations in the HDAC2 and Tip60 binding pattern in the APP larval heads (Supplemental Fig. 4A-D). Taken together, these results suggest that Tip60 exerts its neuroprotective action at least in part *via* protection against inappropriate repressor HDAC2 genome-wide enrichment along neuronal gene bodies.

**Figure 5:**
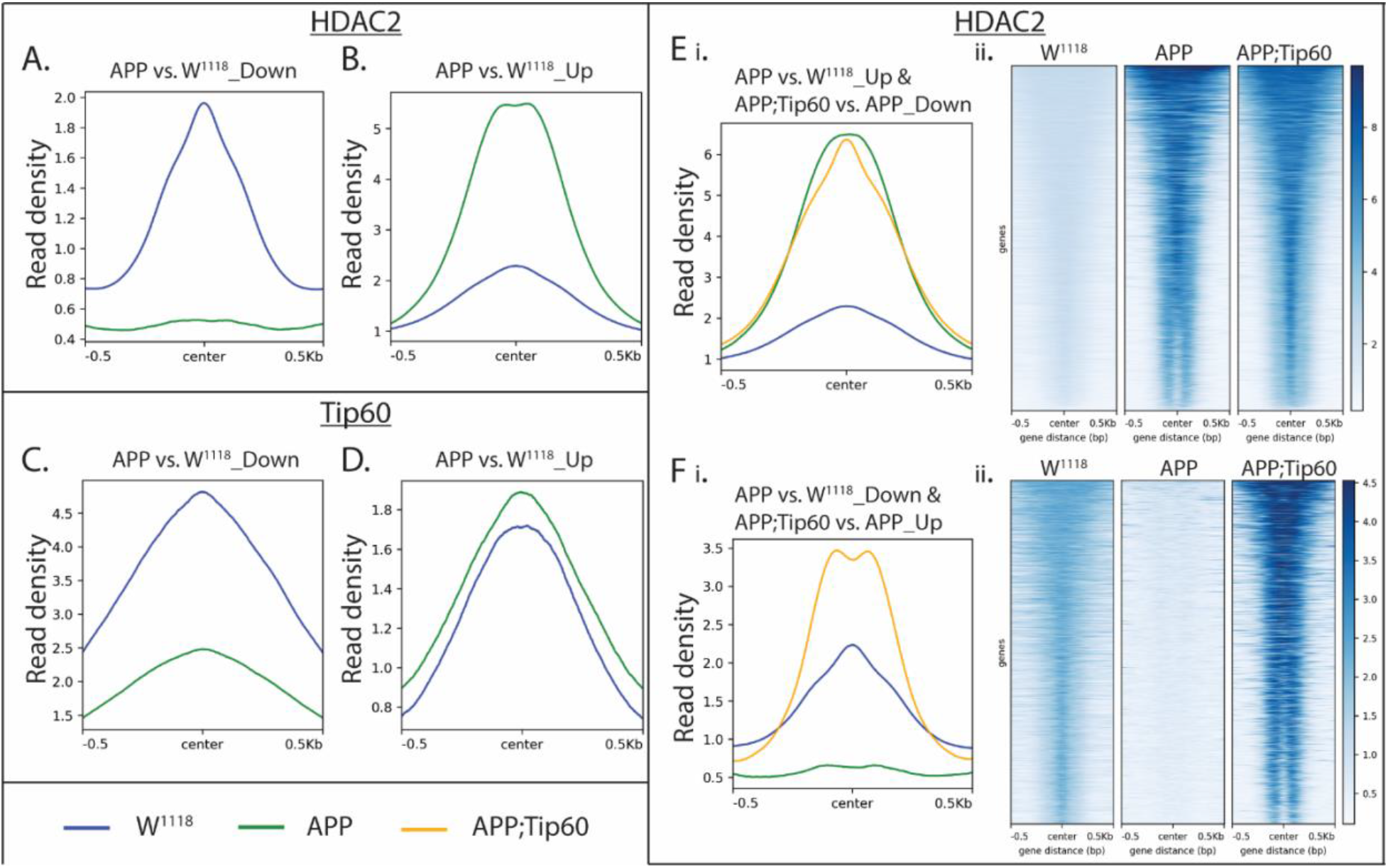
Tip60 expression protected against alterations in the HDAC2 binding pattern along the gene body in APP larval heads. (A & B) Profile plots representing decreased (A) and increased (B) binding of HDAC2 in APP larval heads. (C & D) Profile plots representing the decreased (C) and increased (D) binding of Tip60 in APP larval heads. Profile plots also represent the significant increase in HDAC2 binding (B) and decrease in Tip60 binding (C) in APP larval heads. (E i. & F i.) Profile plots representing the rescue effect (reversal in APP-induced binding pattern) of Tip60 expression on HDAC2 binding. (E ii. & F ii.) The corresponding heatmaps represent the Tip60 rescue effect. Sequencing data centered + /− 0.5 kilobase from the center region of the gene body.

### Tip60/HDAC2 co-regulated genes functionally modulate AD neurodegeneration *in vivo* and are conserved in the human AD brain

We observed that Tip60 and HDAC2 are recruited to genes in a binding enrichment pattern that is disrupted during early AD neurodegeneration, predominantly by HDAC2 binding over Tip60. To identify genes associated with Tip60 and HDAC2 that are misregulated under early AD-associated APP conditions, we compared all differentially expressed genes from our RNA-seq analysis (APP vs. w118 and APP vs. APP;Tip60) with the protein-encoding genes bound by Tip60 in control w^1118^ larval heads and the protein-encoding genes bound by HDAC2 binding in APP larval heads (Supplemental Fig. 5A). Remarkably, this analysis revealed that 77% (or 3137 genes) of the total number of genes identified were identical direct target genes for both HDAC2 and Tip60. These results indicate that Tip60 and HDAC2 co-regulate an identical set of genes and that this control is altered during early AD conditions at least in part by enhanced HDAC2 binding that may also displace Tip60 binding. Comparison of the top twenty (20) biological processes, enriched in gene ontology analysis, each for Tip60 and HDAC2 protein-encoding genes revealed that 17 of these biological processes are identical, further confirming that Tip60 and HDAC2 co-regulate overlapping biological processes (Supplemental Fig. 5B & 5C). These processes included axon guidance, associative learning, and neuron differentiation, underscoring the importance of the co-regulatory function of Tip60 and HDAC2 in neural function and cognition.

We next asked whether these genes are functionally involved in modifying AD-associated neurodegeneration, *in vivo*. To address this question in an unbiased fashion, we selected 50 genes from the top 20 enriched biological processes that were present in both Tip60 and HDAC2 GO analysis (Supplemental Table 5). To assess whether these 50 genes could functionally modulate AD neurodegeneration *in vivo* we used the well-characterized *Drosophila* eye screen that enabled us to assess a gene ability to functionally modulate human tau-driven AD-associated neurodegeneration. To this end, the GMR-Gal4 driver was used to drive the expression of the mutant form of human tau V337M in all retinal cell types. Expression of h-tauV337M in the retina causes a moderately rough eye phenotype at 25°C, characterized by fused and disordered ommatidia with missing mechanosensory bristles (Blard et al., 2007). We determined whether RNAi-mediated knockdown of the genes of interest was able to modify this Tau-induced phenotype by comparing the rough eye phenotype of the Gal4-GMR Tau control flies to the surface of the control Gal4-GMR Tau crossed with RNAi flies. We found that out of 38 genes we were able to obtain RNAi fly lines for, 14 genes showed either enhancement or suppressing of the GMR Tau rough eye phenotype. The functions of these 14 genes include diverse roles in neuronal function and neurodegenerative disease and are referred to here as “Tip60/HDAC2 AD genes”: Shroom, oc, nwk, nmo, Syn, Appl, Dop1R1, RhoGAP100F, NetB, flw, trx, Thor, Dl, & CG7275. The results of the GMR Tau eye screen functionally triaged our mass data sets from both ChIP and RNA sequencing to further streamline mechanistic analysis underlying Tip60 and HDAC2 co-regulation of genes functionally involved in early AD-associated neurodegeneration.

We found that increased Tip60 protects against inappropriate genome-wide enhanced HDAC2 enrichment along the neuronal gene bodies during early AD linked neurodegeneration in APP larval heads. To expand these findings at high resolution, we mapped binding enrichment of both Tip60 and HDAC2 in APP, APP;Tip60, and w^1118^ larval heads along the 14 Tip60/HDAC2 AD gene loci we had identified (Fig. 6 and Supplemental Fig. 6). These genes regulate roles in synaptic plasticity and neuronal developmental processes that include synaptic vesicle function (Syn & nwk), axonal outgrowth (NetB), neuronal signaling pathways (nmo, flw, Dop1R1, Dl, Appl, & RhoGAP100F), gene regulation (trx & Thor), actin filament formation and stabilization (Shroom), and neurodevelopment process (oc) (Larkin et al., 2020). Tip60 and HDAC2 not only both bind within each of these genes in all fly genotypes analyzed (APP, APP;Tip60, w^1118^) but remarkably, at almost identical genomic coordinates, suggesting that Tip60 and HDAC2 are co-recruited to the same docking sites within gene loci. Further, the same trend of inappropriate enhanced HDAC2 enrichment in APP vs. w^1118^ control that was protected against upon increased Tip60 levels (APP;Tip60 vs. APP) was observed at almost all of these genomic coordinates (Supplemental Table 6). Taken together our results support a model by which Tip60 and HDAC2 co-regulate neuronal target genes *via* recruitment to overlapping binding sites within gene bodies.

**Figure 6:**
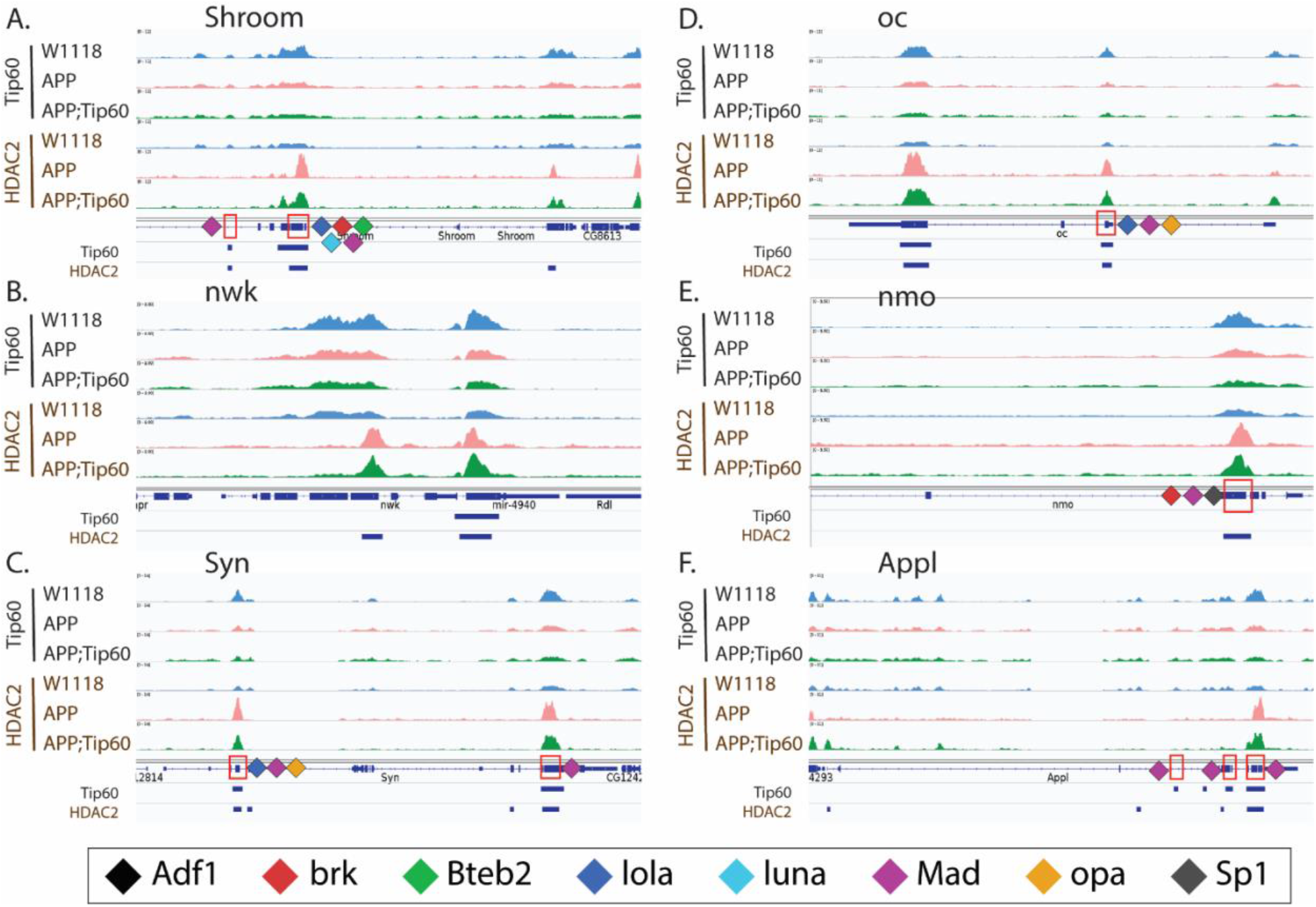
Tip60 and HDAC2 bind at similar genomic coordinates and co-regulate synaptic plasticity and neuronal developmental process-related genes. (A-F) Genome browser track view of Tip60 and HDAC2 peaks in three genotypes (w1118, APP, and APP;Tip60) at the Shroom (A), nwk (B), Syn (C), oc (D), nmo (E), and Appl (F) loci. Below the tracks, the gene features panel has loci marked: representing the transcription factor (Adf1, brk, Bteb2, lola, luna, Mad, opa, and Sp1) binding sites. The blue bars below the gene features panel depicts the regions bound by Tip60 and HDAC2. These genes with significantly enriched peaks exhibit a prominent phenotypical difference in eye screen.

Tip60 and HDAC2 both can interact with transcription factors (TFs) that aid in their gene recruitment and regulatory functions (Aghdassi et al., 2012; Frank et al., 2003; Hlubek et al., 2001; Tea, Chihara, & Luo, 2010; Yang, Inouye, Zeng, Bearss, & Seto, 1996) that we speculate are disrupted in early AD stages. Thus, we asked whether there are conserved TF motif binding sites within genes altered for both expression and Tip60 and HDAC2 binding during early AD stages. To address this question, genes selected for this analysis were triaged by comparing ChIP-Seq and RNA-Seq data sets for AD-associated alterations (APP vs. w^1118^) to select for down-regulated genes with reduced Tip60 and enhanced HDAC2 binding and up-regulated genes with enhanced Tip60 and reduced HDAC2 binding. Only those gene alterations that were protected against by increased Tip60 levels were selected for motif analysis. The selected genes were termed as up-regulated rescue (UpRegRes) list and a down-regulated rescue (DownRegRes) list (Supplemental Fig. 7). GO analysis of these genes revealed that the top biological processes were enriched for functions in learning & memory, axon guidance & extension, neurogenesis & neuron development, and gene silencing & chromatin modification (Supplemental Table 7), further underscoring the importance of Tip60 and HDAC2 in neuronal functions disrupted in AD. Motif enrichment analysis was performed to identify the TFs controlling the rescue genes’ transcription. With HDAC2 encompassing the rescue list, Tip60 bound coordinates altered by APP expression were also included for motif discovery (Supplemental Table 8). The analysis revealed eight TFs with neuronal functions and gene control (Fig. 7A) and motif regions within the Tip60/HDAC2 AD genes (Fig. 7B). Remarkably, many of these TFs are located either within or close to the identified Tip60 and HDAC2 co-peaks (Fig. 6 and Supplemental Fig. 6). Notably, 9 of the 14 Tip60/HDAC2 AD genes have Mad binding sites. Our results suggest that recruitment of Tip60 and HDAC2 by common TFs within gene bodies may be a general mechanism by which these chromatin regulators co-regulate neuronal gene expression.

**Figure 7:**
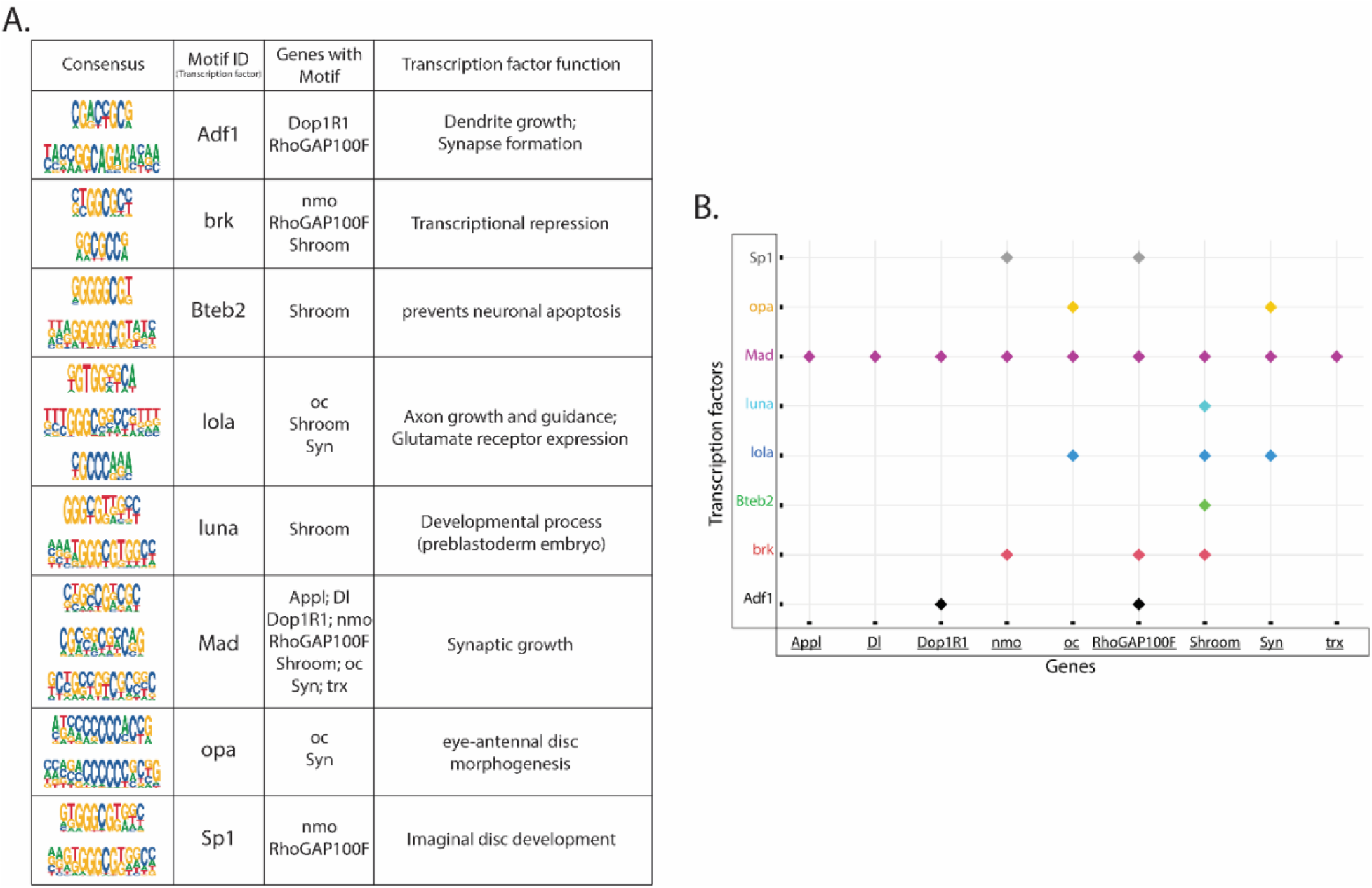
Transcription factor (TF) motifs significantly enriched within the rescue gene list and the associated Tip60/HDAC2 AD genes. TF motifs were identified using the MEME-Chip platform (CentriMo). (A) Consensus sequences and their corresponding TFs bound and the associated Tip60/HDAC2 AD genes. (B) Plot representing the association of Tip60/HDAC2 AD genes and the TFs.

Finally, we asked whether the Tip60/HDAC2 binding alterations and gene dysregulation we observed in APP larval heads were also reflected at the protein level. To address this, we used mass spectrometry (MS) analysis of proteins isolated from the larval brains of w^1118^, APP, and APP;Tip60 genotypes to identify significantly differentially regulated proteins [abs(FC) > 1.5 & q-value <= 0.1] with APP (APP vs. w^1118^) and Tip60 (APP;Tip60 vs. APP) expression (Supplemental Table 9). Analysis of the ~1100 most enriched proteins, identified by MS, revealed that 74 of these proteins were altered in their levels in the APP larval brain and 67 of these in Tip60 expressed brains (Supplemental Fig. 8). Gene ontology analysis revealed that these proteins regulate methylation [histone (Art1 & Art4) & mRNA (Art4)], axon guidance & transport (Dys), nucleocytoplasmic shuttling (Ntf-2), and glutamate (Galphas) & cholinergic (Dys) pathways (Supplemental Table 10). Comparison of proteomics and next-generation sequencing (RNA-seq & ChIP-seq) data reveal that 23% (17/74) of these altered proteins are directly encoded by Tip60/HDAC2 co-regulated genes misregulated in the APP larval brain and 11% (7/67) in the Tip60 expressed brains. (Fig. 8 and Supplemental Table 11). These results suggest that early AD-associated alterations in epigenetic gene regulation persist to the protein level.

**Figure 8:**
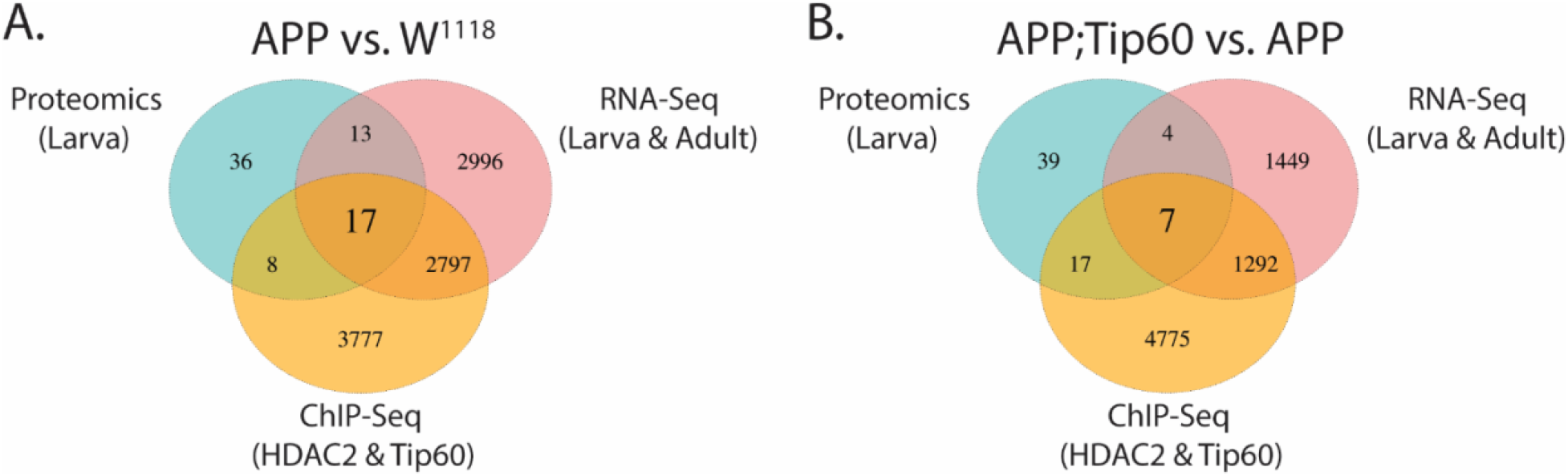
RNA-seq, ChIP-seq, and mass spectrometry data convey the integrative and independent gene expression regulation induced by APP and Tip60 expression. (A & B) Venn diagram of differentially regulated genes in the third instar larval and adult heads (RNA-seq), genes with differentially binding of Tip60 and HDAC2 in the third instar larval heads (ChIP-seq), and differentially regulated proteins in the third instar larval heads (mass spectrometry) from (A) APP vs. w1118 comparison and (B) APP;Tip60 vs. APP comparison.

### Tip60/HDAC2 co-regulation of neuronal genes is disrupted in hippocampus of AD patients

Neuronal gene co-regulation by antagonizing epigenetic enzymes in the human brain has not been investigated previously. A subset of Tip60 and HDAC2 co-regulated direct target genes we identified from our *Drosophila* ChIP-Seq and RNA-Seq analysis that also modify Tau pathology have human orthologs. To confirm human AD disease relevance, we asked whether these same human orthologs are also co-targets of Tip60 and HDAC2 in the human hippocampus and are epigenetically misregulated in the hippocampus from AD patients as we observed in the AD-associated APP fly model. To address these questions, we performed ChIP analysis using chromatin prepared from age-matched human healthy control and AD hippocampus. We quantified enrichment of Tip60 and HDAC2 within gene bodies using real-time PCR. Remarkably, all 14 genes tested were found to be direct gene targets for both Tip60 and HDAC2 in the human hippocampus (Fig. 9). Further, ChIP analysis using chromatin from AD patients revealed that Tip60 enrichment was significantly decreased at 12 of the 14 genes (Fig. 9A). Further, HDAC enrichment was also altered with an increase at 7 of the 14 genes tested and a decrease at 5 of the 14 genes tested (Fig. 9B). Remarkably, five of these genes (Fchsd2, otx1, syde1, ppp1cb, shroom2, dcaf13, and nlk) showed opposite trends in Tip60/HDAC2 binding in the human AD hippocampus, similar to what we observed in AD larval heads. Our findings reveal that Tip60/HDAC2 co-regulatory mechanisms underlying neuronal gene expression that are disrupted during early AD stages in the fly brain and protected against by increased Tip60 are conserved in the hippocampus of human AD patients.

**Figure 9.**
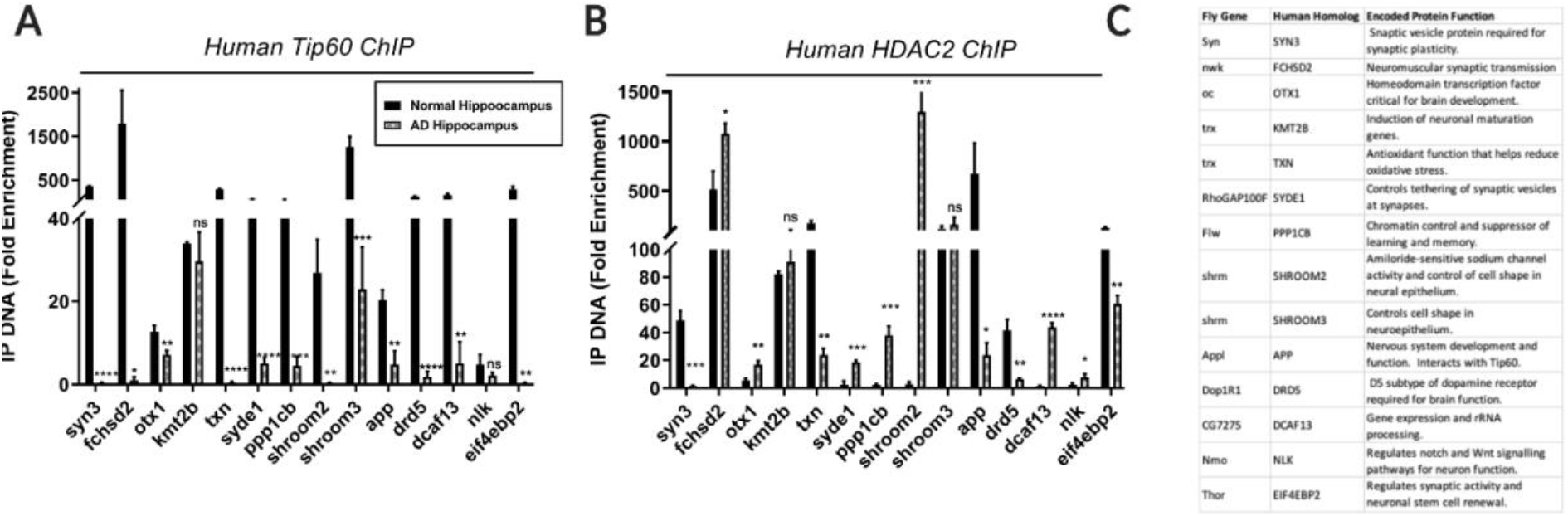
Human homologs of co-Tip60/HDAC2 Drosophila neural gene targets exhibit conserved Tip60 and HDAC2 binding patterns in normal versus AD patient hippocampi. Chromatin was isolated from healthy control and AD hippocampus (n=3 brains per condition). Histograms represent ChIP enrichment using antibodies to (A) Tip60 and (B) and HDAC2. All data are from three independent experiments. Statistical significance was calculated using unpaired Student’s t test. *p < 0.05, **p < 0.01, ***p < 0.001, ****p < 0.0001. Error bars indicate SEM. (See Figure X-1 for primer sequences) (C) Table depicting Drosophila and human homolog gene names and conserved gene functions.

**Figure 10.**
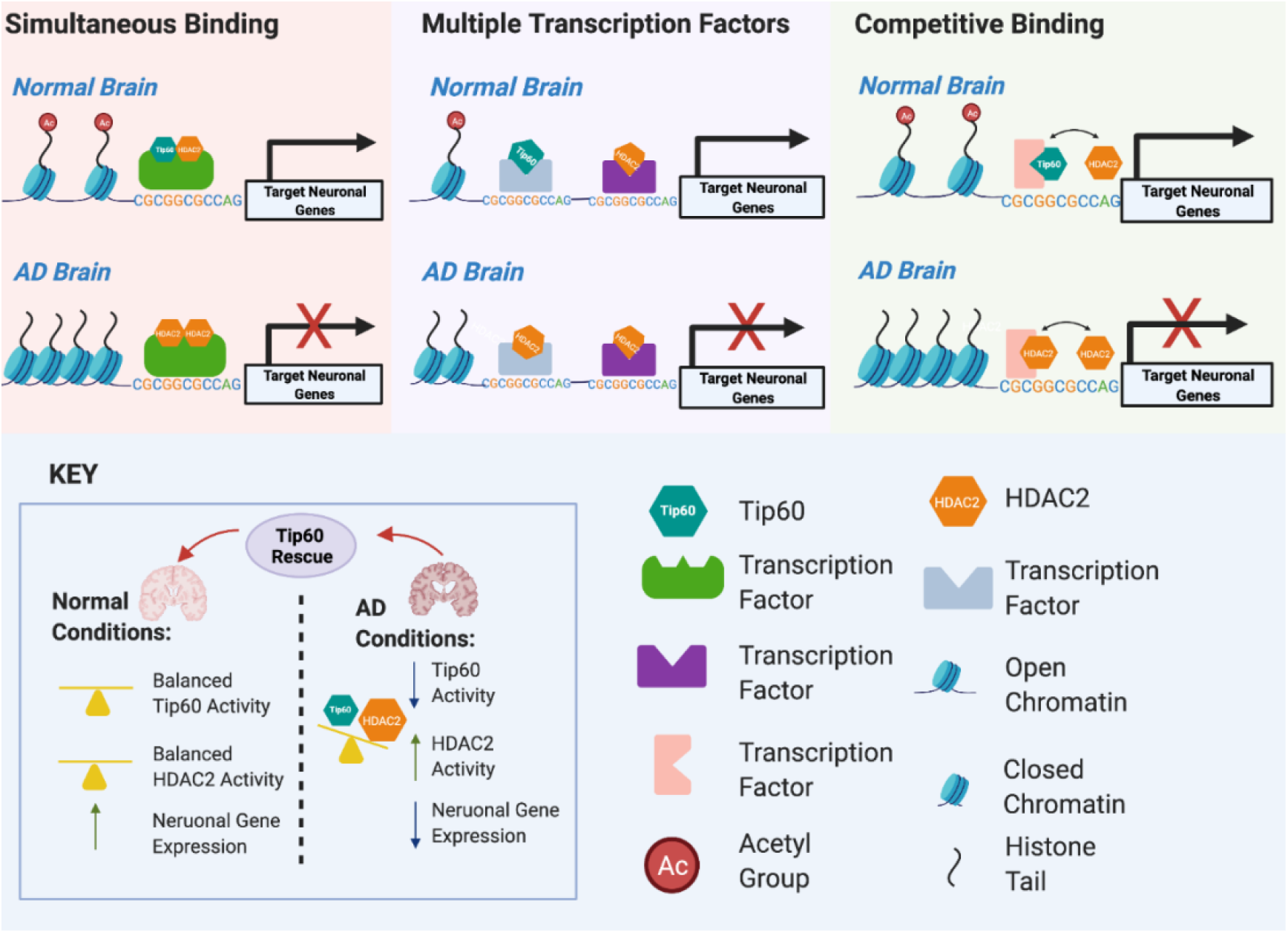
Model for Tip60 and HDAC2 co-mediated neuronal gene control. Our results support a model by which transcription factors (TFs) within a given neuronal gene body serve as docking sites for recruitment of both HDAC2 and Tip60 either simultaneously to the same TF, separately to multiple TFs within close proximity to one another or competitively to a given TF. We speculate that these scenarios are not mutually exclusive of one another and may explain the rapid histone acetylation changes within activity-dependent neural genes that drive their swiftly fluctuating transcriptional responses. Early disruption of Tip60/HDAC homeostasis in AD causes enhanced HDAC2 recruitment with concomitant gene disruption. Increasing Tip60 protects against altered HAT/HDAC homeostasis in the brain to maintain appropriate neuronal gene expression profiles and neural health.

## DISCUSSION

Here we report the first genome-wide HAT versus HDAC profiling study assessing epigenetic alterations initiated during early stages of AD-associated neurodegeneration modeled in the *Drosophila* APP larval brain. A key finding from our analysis revealed that both Tip60 and HDAC2 binding is not exclusively restricted to promoter regions but also enriched predominantly along the gene bodies, suggesting these enzymes may act to both initiate and then maintain gene regulatory control in a poised state (Greer et al., 2015; L. Wang et al., 2017; Z. Wang et al., 2009). Additionally, since gene-body bound TFs also regulate RNA splicing by binding to pre-mRNAs to recruit HATs that increase histone acetylation to facilitate RNA Polymerase elongation and exon exclusion or HDACs that reduce histone acetylation to slow RNA Polymerase elongation and exon inclusion, Tip60 and HDAC2 might also function to regulate RNA splicing of target genes (Greer et al., 2015; Rambout, Dequiedt, & Maquat, 2018). Further, we observed robust alterations in binding enrichment for both HDAC2 and Tip60 in the AD larval brain well before amyloid plaque accumulation and lethality, indicating that chromatin remodeling changes are an initial event in neurodegenerative progression and not a consequence. Notably, our analysis showed a predominant increase in binding of HDAC2 (Fig. 5B) and a decrease in binding of Tip60 (Fig. 5C) within central gene bodies of their target loci. These findings expand prior studies showing enhanced HDAC2 recruitment to a focused subset of synaptic genes in AD fly (Panikker et al., 2018) and mouse models (Graff et al., 2012) by revealing for the first time that an increase in HDAC2 binding is a broad genome-wide AD-associated phenomenon that occurs significantly within gene bodies resulting in their dysregulation. A similar complimentary trend in a marked reduction in genome-wide H4K16 acetylation in the human AD brain (Nativio, Donahue, Berson, Lan, Amlie-Wolf, Tuzer, Toledo, Gosai, Gregory, Torres, et al., 2018), which notably is the preferential acetylation target for Tip60, has recently been reported. Thus, our results indicate that some histone acetylation changes (X. Lu, Wang, Yu, Yu, & Yu, 2015; Nativio, Donahue, Berson, Lan, Amlie-Wolf, Tuzer, Toledo, Gosai, Gregory, Torres, et al., 2018; Stilling & Fischer, 2011) functionally contributing to AD may be initiated at the level of altered Tip60 and HDAC2 antagonistic enzyme recruitment within the central gene body regions.

Another significant finding originating from our work is that Tip60 and HDAC2 co-regulate a similar set of genes that function in cognition linked neural processes disrupted early in AD progression. Comparison of enriched HDAC2 gene targets in the APP larval heads with Tip60 gene targets in the w^1118^ larval heads revealed that, remarkably, 77% of these genes are identical and misregulated in the AD fly brain (Supplemental Fig. 5A). Further, gene ontology analysis of Tip60 versus HDAC2 target genes revealed that 17 of the top 20 most enriched biological processes identified for each enzyme also overlapped and included functions like axonal guidance, associative learning, and neuron differentiation: underscoring their importance in cognitive function and relevance to AD (Supplemental Fig. 5B & 5C). Thus, while other groups have proposed that HAT and HDAC enzymatic activities may both be present in close proximity to each other on gene regulatory regions (Peserico & Simone, 2011; Yamagoe et al., 2003) we are the first to report co-docking of Tip60/HDAC2 on chromatin targets that mediates a co-regulatory function for neural genes at a genome-wide level. Finally, we find that almost one-fourth of the proteins altered in the APP larval brain (17/74) are encoded by dysregulated Tip60/HDAC2 co-target genes (Fig. 8A), indicating that such early AD-associated Tip60/HDAC2 epigenetic alterations persists at the RNA and the protein level.

How might Tip60 and HDAC2 be co-recruited to similar genomic loci within neural genes? It is well-documented that both HATs and HDACs interact with the same TF that facilitates their recruitment to gene loci to promote chromatin remodeling and transcriptional control. For instance, NF-kB interacts with and is acetylated by p300/CBP and deacetylated by HDAC1/HDAC2 to increase and decrease target gene expression, respectively (Chen & Greene, 2004). However, whether HATs and HDACs can bind simultaneously to the same gene by being recruited by either different TFs in close proximity within a given gene locus or by the same TF remains to-be elucidated. Here, in our motif Enrichment Analysis of Tip60 and HDAC2 ChIP-Seq peaks, we identify eight TFs with known neuronal functions and gene control (Fig. 7) that are located either within or in proximity to the Tip60 and HDAC2 co-peaks we identified within AD-associated neural gene loci (Fig. 6 and Supplemental Fig. 6). These findings indicate that these TFs are involved in the co-recruitment of Tip60 and HDAC2 to common gene regulatory regions. Most notably, Mad binds to 9 of the 14 AD-associated genes we analyzed and, remarkably, is present at the identical coordinates within co-Tip60 and HDAC2 peaks within Appl (β amyloid protein precursor-like), shroom, oc, synapsin and delta genes (Fig. 6, Supp. Fig. 6). Accordingly, in prior studies, Mad has been shown to interact with both Tip60 and HDAC2 in other systems to activate and repress gene expression, respectively (Frank et al., 2003; Laherty et al., 1997). Our results support a model by which Mad, along with other TFs within a given gene body, serve as docking sites for recruitment of both HDAC2 and Tip60 either separately and within proximity to one another or simultaneously, thus keeping genes poised for rapid activation or repression. We speculate that these scenarios are not mutually exclusive of one another and, importantly, may explain the rapid histone acetylation changes within activity-dependent neural genes that drive their swiftly fluctuating transcriptional responses (Karnay & Elefant, 2017; Katan-Khaykovich & Struhl, 2002; Peserico & Simone, 2011). Intriguingly, some of the TFs we identify have been previously implicated in AD. For example, Sp1 dysregulation identified in the AD frontal cortex has been proposed to alter its regulation of APP and Tau target genes (Citron, Dennis, Zeitlin, & Echeverria, 2008), while human SMAD (human ortholog of fly Mad) activity is also reduced in the AD brain, causing dysregulation of downstream signaling pathway mediated gene expression (Ueberham et al., 2014).

In the present study, a pivotal discovery with clinical relevance is that increased Tip60 levels protect against altered HDAC2 binding and restoration of appropriate gene expression in the larval brains. Essentially, such Tip60 mediated neuroprotection against epigenetic gene dysregulation is a genome-wide phenomenon as evidenced by our observation that 5400 genes display inappropriate enhanced HDAC2 binding and that increased Tip60 protects against such increases for 74% (3981/5400) of these affected genes in the AD larval brain (Figure 4B). Interestingly, we observed such inappropriate enhanced HDAC2 binding significantly in the gene body’s central region (Figure 5). Further, high-resolution mapping of Tip60 and HDAC2 peaks within AD-associated neuronal genes reveal that enhanced HDAC2 and reduced Tip60 binding in the APP larval head occurs within several Tip60/HDAC2 co-docking sites, with such inappropriate enhanced HDAC2 enrichment reduced with increased Tip60 levels (Fig. 6, Supplemental Fig. 6, and Fig. 5E). Similar trends in altered Tip60/HDAC2 co-regulation of human orthologs of these genes were observed in the human AD hippocampus (Fig. 9), highlighting human relevance and the remarkable conservation in Tip60/HDAC2 epigenetic mechanisms between AD flies and human patients. Together, our findings support a model that increased HDAC2 in the AD larval and human brain (Graff et al., 2012) displaces genome-wide Tip60 recruitment within gene bodies that may be initiated at co-Tip60/HDAC2 docking sites, causing harmful changes in gene expression that persist and worsen during disease progression. Tip60 may mediate its neuroprotective role in epigenetic gene control by either reducing HDAC2 levels, a phenomenon which we previously demonstrated to occur at the transcriptional level (Panikker et al., 2018) and/or by displacing inappropriate enhanced HDAC2 binding levels to restore Tip60 mediated gene regulation.

Our study proposes a mechanism involving aberrant Tip60 and HDAC2 co-recruitment to genes genome-wide to explain how histone acetylation changes are initiated in AD, providing informative directions for chromatin-mediated therapeutic avenues. For example, HDAC inhibitors (HDACi) lack target specificity and act to increase global acetylation (Fischer, Sananbenesi, Mungenast, & Tsai, 2010; Haberland, Montgomery, & Olson, 2009; Johnson et al., 2013), reducing their applicability as safe cognition promoting therapeutics, thus promoting exploration into more specific HAT activators that can potentially reset AD associated site specific histone aceytlation disruption. Our findings underscore this concept by showing that HDAC2 has reduced gene target specificity compared with Tip60, as evidenced by the far more HDAC2 genome-wide target genes altered in the APP larval brain (Fig. 4A) when compared to Tip60. Nevertheless, increased Tip60 specifically protects against altered HDAC2 binding at most genes in the APP larval heads (Fig. 4B) and at many of the co-Tip60/HDAC2 docking sites with TF binding motifs (Fig. 6, Supplemental Fig. 6, and Fig. 7), highlighting the relevance for Tip60 and/or Tip60/HDAC2 interacting TFs as more specific therapeutic targets. Further, these Tip60/HDAC2 binding alterations, at specific gene loci, before Aβ accumulation is detectable, support these sites as potentially valuable early AD biomarker “hot spots” that are easy to track. Recently, we reported that disruption of Tip60 and HDAC2 balance in the brain is a common event in other neurodegenerative diseases modeled in *Drosophila*: HD, ALS, and PD (Beaver et al., 2020). Further studies may reveal a therapeutic potential for targeting Tip60 in these disorders as well. Together, our findings warrant future epigenetic therapeutic studies intended to restore Tip60 mediated histone acetylation homeostasis for earlier and more selective treatment for AD and potentially other neurodegenerative disorders.

## MATERIALS AND METHODS

### Fly stocks

Fly strains and crosses. All fly lines were raised under standard conditions at 22°C on standard yeasted Drosophila media (Applied Scientific Jazz Mix Drosophila Food; Thermo Fisher Scientific, MA, USA). The pan-neuronal driver elav C155 and the transgenic UAS lines carrying human APP 695 isoform (UAS-APP) were obtained from Bloomington Drosophila Stock. Generation and characterization of double-transgenic UAS APP;Tip60 WT fly lines are described in Pirooznia et al. (2012). The w^1118^ line served as the genetic background control. All experimental crosses were performed at normal physiological temperature of 25°C with 12 hour light/dark cycles.

### Immunofluorescence, imaging, and quantification

For anti-Aβ42 immunofluorescence samples were prepared as described in Zhang et al., (2020). Briefly, larval or adult brains were dissected in PBS, fixed in fixation buffer containing 0.7% paraformaldehyde and 0.9% lysine for 1 h at room temperature, washed three times in PBS containing 0.5% Triton X-100 (PBST) for 15 min each time at room temperature, and blocked for 1 h at room temperature in PBST containing 5% normal goat serum, and incubated with primary anti-Aβ42 (1:100, #05-831-I, Millipore, MA, USA) antibody in blocking solution overnight at 4°C. Samples were washed three times in PBST for 15 min each time at room temperature and incubated with goat anti-mouse Alexa Fluor 488 (1:300, #A28175, Invitrogen, CA, USA) and propidium iodide (PI, a final concentration of 1.5 μM) for 2 h at room temperature. After washing three times in PBST for 15 min each time, samples were mounted in VECTASHIELD antifade mounting media (Vector Laboratories, CA, USA).

For imaging, samples were analyzed as described in Zhang et al., (2020). Confocal microscopy was performed using a ZEISS microscope (LSM 700, ZEISS, NY, USA). The optical intervals were 5.94 μm z-sections for 100× magnifications and 0.79 μm z-sections for 200× magnifications. The optical intervals were determined by the optimized pinhole diameters which are 33.3 μm at 1 Airy Unit (AU) for 100× magnification and 25.1 μm at 1 AU for 200× magnification. Consecutive z-stacks through the entire Kn were used for quantification. Consecutive subsets of the z-stacks approximately at the level of center Kn were used for the final projection and display. The quantification of Aβ plaques and apoptosis in different genotypes was measured under 200× magnification using Image J software.

### RNA isolation

Total RNA was isolated from third-instar larval brains or seven-day-old adult heads using the QIAGEN RNeasy Mini Kit (#74106, QIAGEN, MD, USA) following the manufacturer’s protocol. The quality, quantity, and purity of RNA were determined using a Nanodrop spectrophotometer (Thermo Fisher Scientific, MA, USA) and 2100 Bioanalyzer (Agilent Technologies, CA, USA). RNA samples with an RNA integrity number (RIN) ≥ 8.0 were used for sequencing.

### RNA-Seq library preparation, sequencing, and analysis

100 ng of total RNA was used to prepare libraries using TruSeq Stranded Total RNA kit (Illumina, CA, USA) according to the manufacturer’s instructions. The final libraries at the concentration of 4 nM were sequenced on NextSeq 500 platform (Illumina, CA, USA) using 75 bp paired-end sequencing. Raw FASTQ sequencing reads were aligned to the *Drosophila melanogaster* genome (Ensembl version BDGP6) using RNA-Seq by Expectation-Maximization (RSEM) (B. Li & Dewey, 2011). Total read counts were obtained using RSEM’s calculate-expression function. Principal component analysis (PCA) and heatmap clustering (Euclidean distance) were performed to cluster the samples and identify the batch effects and sample heterogeneity. All the plots were constructed using R/Bioconductor.

### Differential gene expression analysis

Differential gene expression between any two genotypes was tested using the DESeq2: a statistical tool that employs shrinkage estimates to compute fold changes (Love, Huber, & Anders, 2014). Raw RNA-Seq read counts from biological replicates of each genotype were used as the input for DESeq2. For both larval and adult data, all three genotypes (w^1118^, APP, and APP;Tip60) were analyzed together using a single model matrix, and the desired pairwise comparisons were then extracted. Only genes that displayed log2FoldChange of ≤ −0.583 and ≥ 0.583 in their expression levels, with an adjusted p-value ≤ 0.05, were used for the UpSet plot (Conway, Lex, & Gehlenborg, 2017) and gene ontology (GO) analysis (FlyEnrichr) (Kuleshov et al., 2016). Among the ontologies in GO analysis, GO Biological Process GeneRIF was included in our downstream analysis. Heatmaps were generated using the ComplexHeatmap package (Gu, Eils, & Schlesner, 2016). The TissueEnrich package is used to calculate enrichment of tissue-specific genes in a set of input genes (Jain & Tuteja, 2019).

### Chromatin immunoprecipitation (ChIP)

Chromatin was extracted and sheared from ~200 third-instar larval heads per replicate. To obtain larval heads, the first 1/3 of the larvae (anterior head region) was isolated. Remaining fat bodies were carefully dissected and discarded. All larval heads were inspected visually to ensure that the entire CNS was intact. Using the GAL4-inducible system to target gene expression exclusively in the nervous system of the larvae ensures virtually no variability in gene expression in the samples used. For IPs, we used truChIP Chromatin Shearing Kit (Covaris Inc., MA, USA) following the manufacturer’s instructions. Briefly, protein–DNA cross-links were made at RT for 5 min with 1% formaldehyde and tissue was pulverized using the CryoPrep (Covaris Inc., MA, USA). Cells were lysed and nuclei were prepared using Covaris lysis buffer. Sonication of DNA was performed using a Covaris E220 Ultrasonicator for 15 min. The sheared chromatin was immunoprecipitated using the EZ-Magna ChIP A Chromatin Immunoprecipitation Kit (Millipore, MA, USA) following the manufacturer’s instructions. Sheering quality and chromatin quantity was determined using Agilent Bioanalyzer DNA 1000 kit (Agilent Technologies, CA, USA). Briefly, ChIP was performed with 30 μg of sheared chromatin using anti-Rpd3 (ab1767, Abcam, MA, USA), anti-Tip60 (ab23886, Abcam, MA, USA). The eluted material from the immunoprecipitation along with an input sample was then purified using a QIAquick PCR purification kit (QIAGEN, MD, USA).

### ChIP-Seq library preparation, sequencing, and analysis

ChIP-Seq libraries were prepared from the ChIP-enriched DNA samples using the Accel-NGS 2SPlus DNA Library Kit (Swift Biosciences, MI, USA), following the 350 base pair insert guide of the protocol. After library preparation, all libraries were normalized and sequenced using the standard Illumina loading protocol on the Illumina NextSeq 500 Sequencer (Illumina, CA, USA). Sequence read fragments were aligned to the *Drosophila melanogaster* BDGP6 genome using the BWA-MEM aligner (H. J. a. p. a. Li, 2013). Samtools was used to filter the resulting alignments to remove reads with mapping quality below q30 and any remaining duplicate reads, and then to merge replicate BAM files for each factor and condition (H. Li et al., 2009). Peak calling was performed on the reads that passed filters for each replicate in addition to the merged alignments using macs2 with default settings (Zhang et al., 2008). The resulting peaks were annotated for genomic features using the HOMER annotatePeaks.pl tool (Heinz et al., 2010). Replicate peak calls were used to estimate the irreproducibility discovery rate (IDR) and create consensus peak sets with IDR ≤ 0.05 (Q. Li, Brown, Huang, & Bickel, 2011). Regions of interest were defined by intersecting the consensus peak sets with Ensembl BDGP6.22 annotation release 98. The featureCounts tool from the subread software package was used to generate read counts for each region of interest (Liao, Smyth, & Shi, 2014). PCA and heatmap clustering (Euclidean distance) were performed to cluster the samples and identify the batch effects and sample heterogeneity. All the plots were constructed using R/Bioconductor.

### Differential binding analysis

Differential binding of peaks (region of interests) between any two genotypes was tested using the DESeq2 (Love et al., 2014). Raw read counts, for each region of interest, from biological replicates of each genotype were used as the input for DESeq2. For Tip60 and HDAC2 samples, all three genotypes (w^1118^, APP, and APP;Tip60) were analyzed together using a single model matrix, and the desired pairwise comparisons were then extracted. Peaks with an adjusted p-value ≤ 0.05 were used for the UpSet plot analysis (Conway et al., 2017). Genes associated with these peaks were further for GO analysis (FlyEnrichr) (Kuleshov et al., 2016). Among the ontologies in GO analysis, GO Biological Process GeneRIF was included in our downstream analysis.

### Visualization of ChIP-seq data

The merged BAM files for each genotype were converted to BPM normalized BigWig files using bamCompare. computeMatrix was used to calculate scores per genome regions (Differentially bound regions from DESeq2) and prepared an intermediate file that can be used with plotHeatmap and plotProfiles (Ramírez et al., 2016). The reference point for the plotting was the center of the region with a window of + /− 0.5 kilobase. For ChIP-Seq track generation, BigWig files were used with Integrated Genomics Viewer (IGV_Linux_2.8.6) (Robinson et al., 2011). The BED files used for IGV contain genomic coordinates of the significantly enriched peaks of genes resulted from the eye-screen (Supplemental Table 6).

### Motif enrichment analysis

We performed DNA motif enrichment analysis, central motif enrichment analysis or CentriMo (Bailey & Machanick, 2012), to detect the positional enrichment of previously characterized TF binding motifs in the Tip60 and HDAC2 bound sequences (Supplemental Fig. 7). The Combined Drosophila Databases (TF motifs) provided in the web version of the CentriMo were used as the input for CentriMo. The default options were used for the analysis, and the statistical significance of discovered motifs was estimated using *P* values and *E*-values derived from a one-tailed binomial test (Supplemental Table 8).

### Protein isolation, identification, and analysis

Protein was extracted from dissected third-instar larval brains of three genotypes (w^1118^, APP, and APP;Tip60) and was sent to Bioproximity LLC for proteomic profiling. Samples were subjected to enzymatic digestion with sequencing-grade trypsin. The digested peptides were cleaned-up by solid-phase extraction (SPE) protocol. Each digestion mixture was analyzed by UPLC-MS/MS (Ultra performance liquid chromatography-tandem mass spectrometer). LC was performed on an Easy-nLC 1200 system (Thermo Fisher Scientific, MA, USA) fitted with a heated, 25 cm Easy-Spray column. The LC was interfaced to a quadrupole-Orbitrap mass spectrometer (Q-Exactive HF-X, Thermo Fisher Scientific, MA, USA). TMGF (Mascot Generic Format) files were searched using X!Tandem and Open Mass Spectrometry Search Algorithm (OMSSA). Protein intensity values were calculated using OpenMS to measure the area under the curve of identified peptides. The Perseus software platform was used for protein quantification, cross-comparisons between genotypes, and multiple-hypothesis testing (Benjamini-Hochberg FDR: t-test p-value adjusted to account for multiple testing) (Tyanova et al., 2016). Proteins with q < 0.05 and |FC| > 1.5, determined as significantly changed proteins, were used for downstream analysis. Protein-protein interaction networks among the significantly changed proteins were visualized using STRING on the Cytoscape platform (Cytoscape_v3.7.2) (Shannon et al., 2003). Functional enrichment analysis was performed using FlyEnrichr (Kuleshov et al., 2016) and GO Biological Process GeneRIF was included in our downstream analysis.

### ChIPqPCR (Human)

For all human studies, human hippocampal samples were obtained from National Disease Research Interchange, with informed consent by all donors. Control brains included three males with an age range of 70 –85 years. AD brains were from one male and two females with an age range of 73-87 years.

Chromatin was extracted and sheared from ~120 mg human hippocampus using truChIP Chromatin Shearing Kit (Covaris Inc., MA, USA) following the manufacturer’s instructions. Briefly, protein–DNA crosslinks were made at RT for 5 min with 1% formaldehyde and tissue was pulverized using the CryoPrep (Covaris Inc., MA, USA). Cells were lysed and nuclei were prepared using Covaris lysis buffer. Sonication of DNA was performed using a Covaris E220 Ultrasonicator for 13 min. The sheared chromatin was immunoprecipitated using the EZ-Magna ChIPA Chromatin Immunoprecipitation Kit (Millipore, MA, USA) following the manufacturer’s instructions. Briefly, ChIP was performed with 50ug of sheared chromatin using anti-Tip60 (ab23886, Abcam, MA, USA), anti-HDAC2 (ab12169, Abcam, MA, USA), and Normal Mouse IgG Polyclonal Antibody control (Millipore, MA, USA). Eluted material from the immunoprecipitation was purified using a QIAquick PCR purification kit (QIAGEN, MD, USA) and used directly for real-time PCR.

qRT-PCRs were performed in a 20 uL reaction volume containing cDNA, 1 M Power SYBR Green PCR Master Mix (Applied Biosystems, CA, USA), and 10 M forward and reverse primers (Supplemental Table 12). Primer sets were designed by NCBI/Primer-BLAST (www.ncbi.nlm.nih.gov/tools/primer-blast/). RT-qPCR was performed using an ABI 7500 Real-Time PCR system (Applied Biosystems, CA, USA) following the manufacturer’s instructions. Fold enrichment for all the respective genes was calculated relative to the non-specific Mouse IgG Polyclonal Antibody control.

### Statistical analysis

Statistical analysis of RNA-Seq, ChIP-Seq, and mass spectrometry (MS) data differences between two groups were considered statistically significant with q < 0.05 (FDR < 0.05, controlled by Benjamini–Hochberg). For ChIP-seq analysis, sample sizes were w^1118^ = 3; APP = 2; APP;Tip60 = 2. For RNA-seq analysis, the sample size for third-instar larva was w^1118^ = 2; APP = 2; APP;Tip60 = 2 and for seven-day-old adult flies was w^1118^ = 3; APP = 3; APP;Tip60 = 3. For MS analysis, sample sizes were w^1118^ = 3; APP = 2; APP;Tip60 = 2.

Model figure created using BioRender.com

## ACKNOWLEDGEMENTS

Research reported in this publication utilized the MetaOmics Shared Resource at Sidney Kimmel Cancer Center at Jefferson Health that was supported by the National Cancer Institute of the National Institutes of Health (NIH) under Award Number P30CA056036. We thank Adam Ertel and Guarav Kumar for their help with the initial bioinformatics analysis. We are grateful for the human subjects and their families for participation in this study. The research was supported by the National Institutes of Neurological Disorders and Stroke of the NIH under Award Number R01HD057939 to F.E.

## References

Aghdassi, A., Sendler, M., Guenther, A., Mayerle, J., Behn, C.-O., Heidecke, C.-D., … Lerch, M. M. (2012). Recruitment of histone deacetylases HDAC1 and HDAC2 by the transcriptional repressor ZEB1 downregulates E-cadherin expression in pancreatic cancer. Gut, 61(3), 439–448.

Bailey, T. L., & Machanick, P. J. N. a. r. (2012). Inferring direct DNA binding from ChIP-seq. 40(17), e128–e128.

Beaver, M., Bhatnagar, A., Panikker, P., Zhang, H., Snook, R., Parmar, V., … Elefant, F. (2020). Disruption of Tip60 HAT mediated neural histone acetylation homeostasis is an early common event in neurodegenerative diseases. Sci Rep, 10(1), 18265. doi:10.1038/s41598-020-75035-3

Berson, A., Nativio, R., Berger, S. L., & Bonini, N. M. (2018). Epigenetic regulation in neurodegenerative diseases. Trends in neurosciences, 41(9), 587–598.

Blard, O., Feuillette, S., Bou, J., Chaumette, B., Frébourg, T., Campion, D., & Lecourtois, M. (2007). Cytoskeleton proteins are modulators of mutant tau-induced neurodegeneration in Drosophila. Hum Mol Genet, 16(5), 555–566. doi:10.1093/hmg/ddm011

Chen, L.-F., & Greene, W. C. (2004). Shaping the nuclear action of NF-κB. Nature Reviews Molecular Cell Biology, 5(5), 392–401. doi:10.1038/nrm1368

Citron, B. A., Dennis, J. S., Zeitlin, R. S., & Echeverria, V. (2008). Transcription factor Sp1 dysregulation in Alzheimer’s disease. Journal of Neuroscience Research, 86(11), 2499–2504. doi:https://doi.org/10.1002/jnr.21695

Conway, J. R., Lex, A., & Gehlenborg, N. J. B. (2017). UpSetR: an R package for the visualization of intersecting sets and their properties. 33(18), 2938–2940.

Fischer, A., Sananbenesi, F., Mungenast, A., & Tsai, L. H. (2010). Targeting the correct HDAC(s) to treat cognitive disorders. Trends Pharmacol Sci, 31(12), 605–617. doi:10.1016/j.tips.2010.09.003

Fossgreen, A., Bruckner, B., Czech, C., Masters, C. L., Beyreuther, K., & Paro, R. (1998). Transgeneic Drosphila expressing human amyloid precursor protein show gamma-secretase activity and a blistered-wing phenotype. Proceedings National Acadamy of Sciences, 96, 13703–13708.

Fossgreen, A., Brückner, B., Czech, C., Masters, C. L., Beyreuther, K., & Paro, R. (1998). Transgenic Drosophila expressing human amyloid precursor protein show gamma-secretase activity and a blistered-wing phenotype. Proc Natl Acad Sci U S A, 95(23), 13703–13708. doi:10.1073/pnas.95.23.13703

Frank, S. R., Parisi, T., Taubert, S., Fernandez, P., Fuchs, M., Chan, H.-M., … Amati, B. (2003). MYC recruits the TIP60 histone acetyltransferase complex to chromatin. EMBO reports, 4(6), 575–580. doi:10.1038/sj.embor.embor861

Graff, J., Rei, D., Guan, J. S., Wang, W. Y., Seo, J., Hennig, K. M., … Tsai, L. H. (2012). An epigenetic blockade of cognitive functions in the neurodegenerating brain. Nature, 483(7388), 222–226. doi:10.1038/nature10849

Greer, C. B., Tanaka, Y., Kim, Y. J., Xie, P., Zhang, M. Q., Park, I.-H., & Kim, T. H. (2015). Histone deacetylases positively regulate transcription through the elongation machinery. Cell reports, 13(7), 1444–1455.

Greeve, I., Kretzschmar, D., Tschape, J.A.., Beyn, A., Brellinger, C., Schweizer, M., Nitsch, T.M., Reifegerste, R. (2004). Age-dependent neurodegenertion and Alzheimer-amyloid plaque formation in transgenic Drosophila. Journal of Neuroscience (16), 3899–38906.

Grothe, M. J., Sepulcre, J., Gonzalez-Escamilla, G., Jelistratova, I., Scholl, M., Hansson, O., … Alzheimer’s Disease Neuroimaging, I. (2018). Molecular properties underlying regional vulnerability to Alzheimer’s disease pathology. Brain, 141(9), 2755–2771. doi:10.1093/brain/awy189

Gu, Z., Eils, R., & Schlesner, M. J. B. (2016). Complex heatmaps reveal patterns and correlations in multidimensional genomic data. 32(18), 2847–2849.

Haberland, M., Montgomery, R. L., & Olson, E. N. (2009). The many roles of histone deacetylases in development and physiology: implications for disease and therapy. Nat Rev Genet, 10(1), 32–42. doi:10.1038/nrg2485

Heinz, S., Benner, C., Spann, N., Bertolino, E., Lin, Y. C., Laslo, P., … Glass, C. K. J. M. c. (2010). Simple combinations of lineage-determining transcription factors prime cis-regulatory elements required for macrophage and B cell identities. 38(4), 576–589.

Hlubek, F., Löhberg, C., Meiler, J., Jung, A., Kirchner, T., & Brabletz, T. (2001). Tip60 is a cell-type-specific transcriptional regulator. The Journal of Biochemistry, 129(4), 635–641.

Iijima, K., Chiang, H. C., Hearn, S. A., Hakker, I., Gatt, A., Shenton, C., … Zhong, Y. (2008). Abeta42 mutants with different aggregation profiles induce distinct pathologies in Drosophila. PLoS One, 3(2), e1703. doi:10.1371/journal.pone.0001703

Iijima, K., Liu, H. P., Chiang, A. S., Hearn, S. A., Konsolaki, M., & Zhong, Y. (2004). Dissecting the pathological effects of human Abeta40 and Abeta42 in Drosophila: a potential model for Alzheimer’s disease. Proc Natl Acad Sci U S A, 101(17), 6623–6628. doi:10.1073/pnas.0400895101

Jain, A., & Tuteja, G. J. B. (2019). TissueEnrich: Tissue-specific gene enrichment analysis. 35(11), 1966–1967.

Johnson, A. A., Sarthi, J., Pirooznia, S. K., Reube, W., & Elefant, F. (2013). Increasing Tip60 HAT levels rescues axonal transport defects and associated behavioral phenotypes in a Drosophila Alzheimer’s disease model. J Neurosci, 33(17), 7535–7547. doi:10.1523/JNEUROSCI.3739-12.2013

Karch, C. M., Cruchaga, C., & Goate, A. M. (2014). Alzheimer’s disease genetics: from the bench to the clinic. Neuron, 83(1), 11–26. doi:10.1016/j.neuron.2014.05.041

Karnay, A. M., & Elefant, F. (2017). Chapter 14 - Drosophila Epigenetics. In T. O. Tollefsbol (Ed.), Handbook of Epigenetics (Second Edition) (pp. 205–229): Academic Press.

Katan-Khaykovich, Y., & Struhl, K. (2002). Dynamics of global histone acetylation and deacetylation in vivo: rapid restoration of normal histone acetylation status upon removal of activators and repressors. Genes & development, 16(6), 743–752. doi:10.1101/gad.967302

Kuleshov, M. V., Jones, M. R., Rouillard, A. D., Fernandez, N. F., Duan, Q., Wang, Z., … Lachmann, A. J. N. a. r. (2016). Enrichr: a comprehensive gene set enrichment analysis web server 2016 update. 44(W1), W90–W97.

Kunkle, B. W., Grenier-Boley, B., Sims, R., Bis, J. C., Damotte, V., Naj, A. C., … Environmental Risk for Alzheimer’s Disease, C. (2019). Genetic meta-analysis of diagnosed Alzheimer’s disease identifies new risk loci and implicates Abeta, tau, immunity and lipid processing. Nat Genet, 51(3), 414–430. doi:10.1038/s41588-019-0358-2

Laherty, C. D., Yang, W.-M., Sun, J.-M., Davie, J. R., Seto, E., & Eisenman, R. N. (1997). Histone Deacetylases Associated with the mSin3 Corepressor Mediate Mad Transcriptional Repression. Cell, 89(3), 349–356. doi:10.1016/S0092-8674(00)80215-9

Larkin, A., Marygold, S. J., Antonazzo, G., Attrill, H., dos Santos, G., Garapati, P. V., … Consortium, F. (2020). FlyBase: updates to the Drosophila melanogaster knowledge base. Nucleic acids research, 49(D1), D899–D907. doi:10.1093/nar/gkaa1026

Li, B., & Dewey, C. N. J. B. b. (2011). RSEM: accurate transcript quantification from RNA-Seq data with or without a reference genome. 12(1), 1–16.

Li, H., Handsaker, B., Wysoker, A., Fennell, T., Ruan, J., Homer, N., … Durbin, R. J. B. (2009). The sequence alignment/map format and SAMtools. 25(16), 2078–2079.

Li, H. J. a. p. a. (2013). Aligning sequence reads, clone sequences and assembly contigs with BWA-MEM.

Li, Q., Brown, J. B., Huang, H., & Bickel, P. J. J. T. a. o. a. s. (2011). Measuring reproducibility of high-throughput experiments. 5(3), 1752–1779.

Liao, Y., Smyth, G. K., & Shi, W. J. B. (2014). featureCounts: an efficient general purpose program for assigning sequence reads to genomic features. 30(7), 923–930.

Love, M. I., Huber, W., & Anders, S. J. G. b. (2014). Moderated estimation of fold change and dispersion for RNA-seq data with DESeq2. 15(12), 1–21.

Lu, X., Wang, L., Yu, C., Yu, D., & Yu, G. (2015). Histone Acetylation Modifiers in the Pathogenesis of Alzheimer’s Disease. Frontiers in cellular neuroscience, 9, 226–226. doi:10.3389/fncel.2015.00226

Lu, X., Wang, L., Yu, C., Yu,D., and Yu, G. (2015). Histone acetylation modifiers in the pathogenesis of Alzheimer’s disease. Frontiers in Cellular Neuroscience, 9(226), 1–8.

Masters, C. L., Bateman, R., Blennow, K., Rowe, C. C., Sperling, R. A., & Cummings, J. L. (2015). Alzheimer’s disease. Nat Rev Dis Primers, 1, 15056. doi:10.1038/nrdp.2015.56

Nativio, R., Donahue, G., Berson, A., Lan, Y., Amlie-Wolf, A., Tuzer, F., … Torres, C. (2018). Dysregulation of the epigenetic landscape of normal aging in Alzheimer’s disease. Nature neuroscience, 21(4), 497.

Nativio, R., Donahue, G., Berson, A., Lan, Y., Amlie-Wolf, A., Tuzer, F., … Berger, S. L. (2018). Dysregulation of the epigenetic landscape of normal aging in Alzheimer’s disease. Nature neuroscience, 21(4), 497–505. doi:10.1038/s41593-018-0101-9

Panikker, P., Xu, S. J., Zhang, H., Sarthi, J., Beaver, M., Sheth, A., … Elefant, F. (2018). Restoring Tip60 HAT/HDAC2 Balance in the Neurodegenerative Brain Relieves Epigenetic Transcriptional Repression and Reinstates Cognition. J Neurosci, 38(19), 4569–4583. doi:10.1523/JNEUROSCI.2840-17.2018

Patel, H., Dobson, R. J. B., & Newhouse, S. J. (2019). A Meta-Analysis of Alzheimer’s Disease Brain Transcriptomic Data. J Alzheimers Dis, 68(4), 1635–1656. doi:10.3233/JAD-181085

Peserico, A., & Simone, C. (2011). Physical and Functional HAT/HDAC Interplay Regulates Protein Acetylation Balance. Journal of Biomedicine and Biotechnology, 2011, 371832. doi:10.1155/2011/371832

Pirooznia, S. K., Sarthi, J., Johnson, A. A., Toth, M. S., Chiu, K., Koduri, S., & Elefant, F. (2012). Tip60 HAT activity mediates APP induced lethality and apoptotic cell death in the CNS of a Drosophila Alzheimer’s disease model. PLoS One, 7(7), e41776. doi:10.1371/journal.pone.0041776

Rambout, X., Dequiedt, F., & Maquat, L. (2018). Beyond Transcription: Roles of Transcription Factors in Pre-mRNA Splicing. Chemical reviews, 118 8, 4339–4364.

Ramírez, F., Ryan, D. P., Grüning, B., Bhardwaj, V., Kilpert, F., Richter, A. S., … Manke, T. J. N. a. r. (2016). deepTools2: a next generation web server for deep-sequencing data analysis. 44(W1), W160–W165.

Robinson, J. T., Thorvaldsdóttir, H., Winckler, W., Guttman, M., Lander, E. S., Getz, G., & Mesirov, J. P. J. N. b. (2011). Integrative genomics viewer. 29(1), 24–26.

Saha, R. N., & Pahan, K. (2006). HATs and HDACs in neurodegeneration: a tale of disconcerted acetylation homeostasis. Cell Death Differ, 13(4), 539–550. doi:10.1038/sj.cdd.4401769

Sanchez-Mut, J. V., & Graff, J. (2015). Epigenetic Alterations in Alzheimer’s Disease. Front Behav Neurosci, 9, 347. doi:10.3389/fnbeh.2015.00347

Shannon, P., Markiel, A., Ozier, O., Baliga, N. S., Wang, J. T., Ramage, D., … Ideker, T. J. G. r. (2003). Cytoscape: a software environment for integrated models of biomolecular interaction networks. 13(11), 2498–2504.

Stilling, R. M., & Fischer, A. (2011). The role of histone acetylation in age-associated memory impairment and Alzheimer’s disease. Neurobiology of learning and memory, 96(1), 19–26. doi:https://doi.org/10.1016/j.nlm.2011.04.002

Tea, J. S., Chihara, T., & Luo, L. (2010). Histone deacetylase Rpd3 regulates olfactory projection neuron dendrite targeting via the transcription factor Prospero. Journal of Neuroscience, 30(29), 9939–9946.

Tyanova, S., Temu, T., Sinitcyn, P., Carlson, A., Hein, M. Y., Geiger, T., … Cox, J. J. N. m. (2016). The Perseus computational platform for comprehensive analysis of (prote) omics data. 13(9), 731.

Ueberham, U., Rohn, S., Ueberham, E., Wodischeck, S., Hilbrich, I., Holzer, M., … Arendt, T. (2014). Pin1 promotes degradation of Smad proteins and their interaction with phosphorylated tau in Alzheimer’s disease. Neuropathol Appl Neurobiol, 40(7), 815–832. doi:10.1111/nan.12163

Wang, L., Zhang, F., Rode, S., Chin, K. K., Ko, E. E., Kim, J., … Qiao, H. (2017). Ethylene induces combinatorial effects of histone H3 acetylation in gene expression in Arabidopsis. BMC genomics, 18(1), 1–13.

Wang, Z., Zang, C., Cui, K., Schones, D. E., Barski, A., Peng, W., & Zhao, K. (2009). Genome-wide Mapping of HATs and HDACs Reveals Distinct Functions in Active and Inactive Genes. Cell, 138(5), 1019–1031. doi:10.1016/j.cell.2009.06.049

Xu, S., Wilf, R., Menon, T., Panikker, P., Sarthi, J., & Elefant, F. (2014). Epigenetic control of learning and memory in Drosophila by Tip60 HAT action. Genetics, 198(4), 1571–1586. doi:10.1534/genetics.114.171660

Yamagoe, S., Kanno, T., Kanno, Y., Sasaki, S., Siegel, R. M., Lenardo, M. J., … Ozato, K. (2003). Interaction of Histone Acetylases and Deacetylases In Vivo. Molecular and cellular biology, 23(3), 1025–1033. doi:10.1128/mcb.23.3.1025-1033.2003

Yang, W. M., Inouye, C., Zeng, Y., Bearss, D., & Seto, E. (1996). Transcriptional repression by YY1 is mediated by interaction with a mammalian homolog of the yeast global regulator RPD3. Proc Natl Acad Sci U S A, 93(23), 12845–12850. doi:10.1073/pnas.93.23.12845

Zhang, Y., Liu, T., Meyer, C. A., Eeckhoute, J., Johnson, D. S., Bernstein, B. E., … Li, W. J. G. b. (2008). Model-based analysis of ChIP-Seq (MACS). 9(9), 1–9.

