## Supplementary material for "Chromatin and transcriptomic profiling uncover dysregulation of the Tip60 HAT/HDAC2 epigenomic landscape in the neurodegenerative brain": S

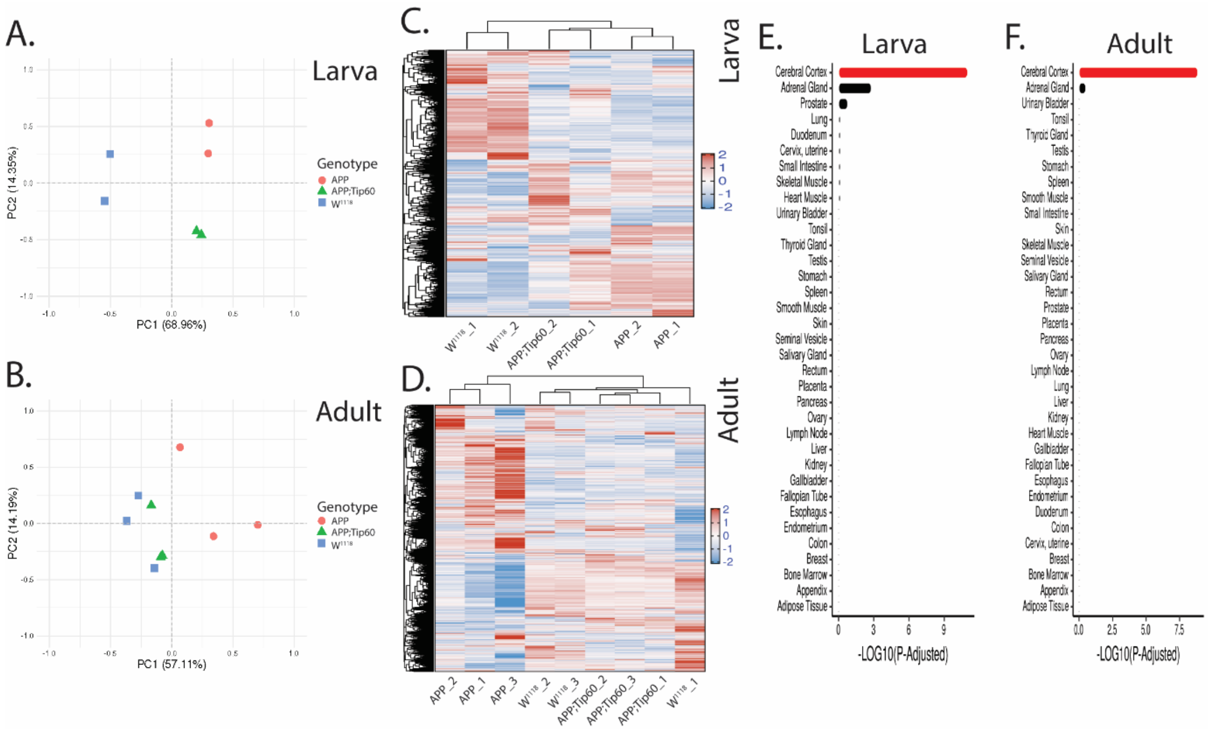


**Figure S1: An overview of the sample clustering and tissue-specific gene set enrichment of RNA-seq data.** Principal component analysis of 3rd-instar larval (A) and adult (B) samples. Samples are color-coded by genotype ($w^{1118}$, APP, and APP;Tip60). Variation among samples is explained by the genotype with distinct clusters in 3rd-instar larval stage (A). Clustering of APP;Tip60 samples with the $w^{1118}$ samples in adult (B) stage reflect the similarity in gene expression profiles between these genotypes. Heatmaps representing the hierarchical clustering of 3rd-instar larval (C) and adult (D) samples of three genotypes. Samples from the same genotype cluster well, particularly in third instar larvae. In both third instar larvae and adults, APP;Tip60 samples are closer to $w^{1118}$ samples indicating the similarity in gene expression profiles between these genotypes. Neural tissue enrichment with the top 2000 genes (human orthologs) regulated (1000 up-regulated and 1000 down-regulated genes) by APP expression (APP vs. $w^{1118}$), in third instar larval (E) and adult (F) heads, reiterates the neural-specific changes with the APP expression and the purity of the sample input for RNA-seq.

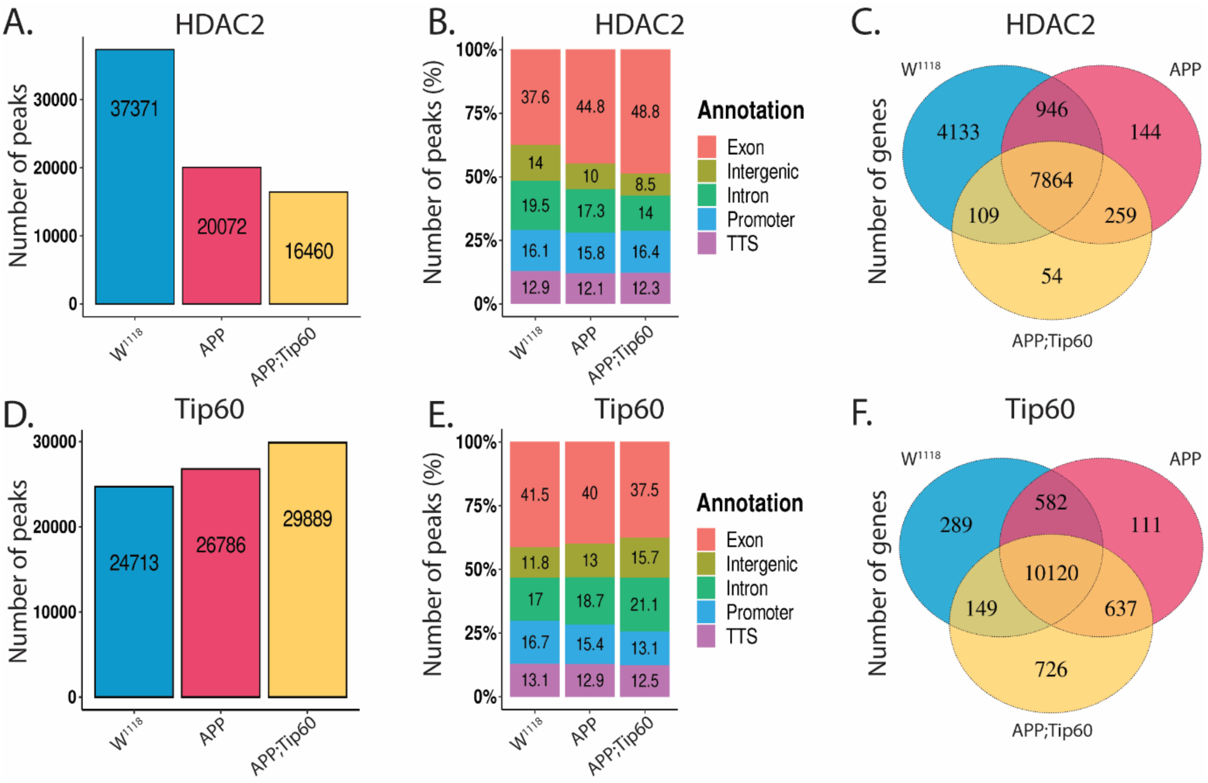


**Figure S2: HDAC2 and Tip60 peaks predominantly enriched over the gene body.** Bar plots comparing the HDAC2 (A) and Tip60 peaks (D) in the third instar larvae, among $w^{1118}$, APP, and APP;Tip60 genotypes. (B & E) The majority of the peaks identified are distributed over exon and intronic regions. Venn diagrams of the genes (peaks bound to) bound by HDAC2 (C) and Tip60 (F) overlap among $w^{1118}$, APP, and APP;Tip60 genotypes with a majority of the genes in common and certain unique genes for each genotype.

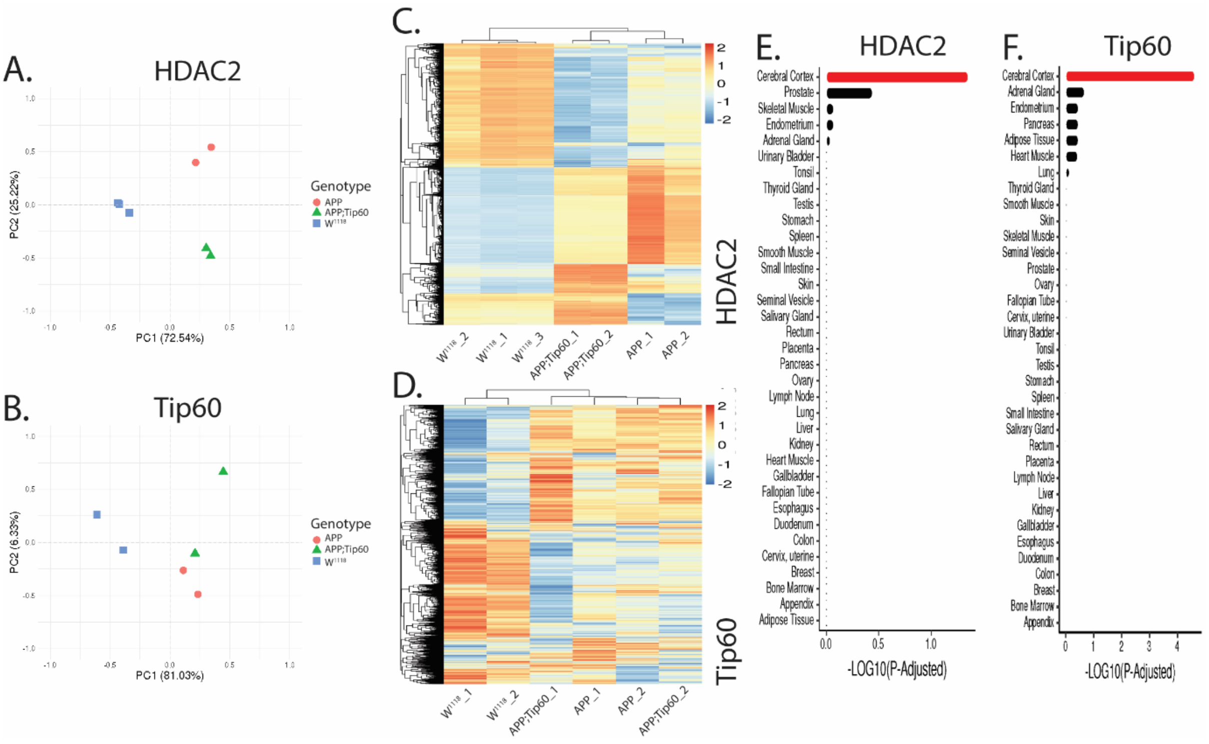


**Figure S3: An overview of the sample clustering and tissue-specific gene set enrichment of ChIP-seq data.** Principal component analysis of the HDAC2 (A) and Tip60 (B) samples. Samples are color-coded by genotype ($w^{1118}$, APP, and APP;Tip60). Variation among samples is explained by the genotype with distinct clusters in HDAC2 (A). Clustering of APP;Tip60 samples with the APP samples, w.r.t. PC1, in both HDAC2 (A) and Tip60 (B) reflects less variation between these genotypes. Nevertheless, these genotypes vary w.r.t. PC2. Heatmaps representing the hierarchical clustering of HDAC2 (C) and Tip60 (D) samples of the three genotypes. Samples from the same genotype cluster well, particularly with HDAC2. In both HDAC2 and Tip60, particularly in HDAC2, the clustering distance between APP;Tip60 and $w^{1118}$ samples is minimal. It indicates a relative similarity in gene expression profiles between these genotypes. Neural tissue enrichment with the top 2000 genes (human orthologs of 1000 increased binding and 1000 decreased binding) differentially bound by HDAC2 (E) and Tip60 (F) by APP expression (APP vs. $w^{1118}$) reiterates the neural-specific changes with the APP expression and the purity of the sample input for ChIP-seq.

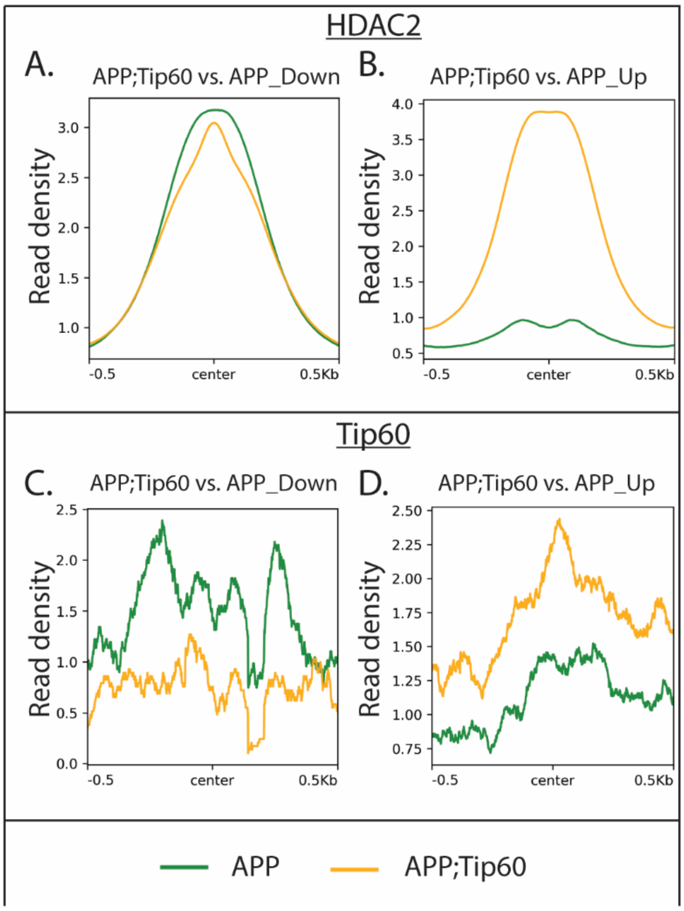


**Figure S4: Tip60 expression reversed the APP-induced HDAC2 and Tip60 binding pattern in third instar larval brain.** (A & B) Profile plots representing the Tip60-induced decreased (A) and increased (B) binding of HDAC2. (C & D) Profile plots representing the Tip60-induced decreased (C) and increased (D) binding of Tip60. (C & D) The roughness in Tip60 binding profile plots is the result of a fewer number of peaks present. Sequencing data centered + /- 0.5 kilobase from the center region of the gene body.

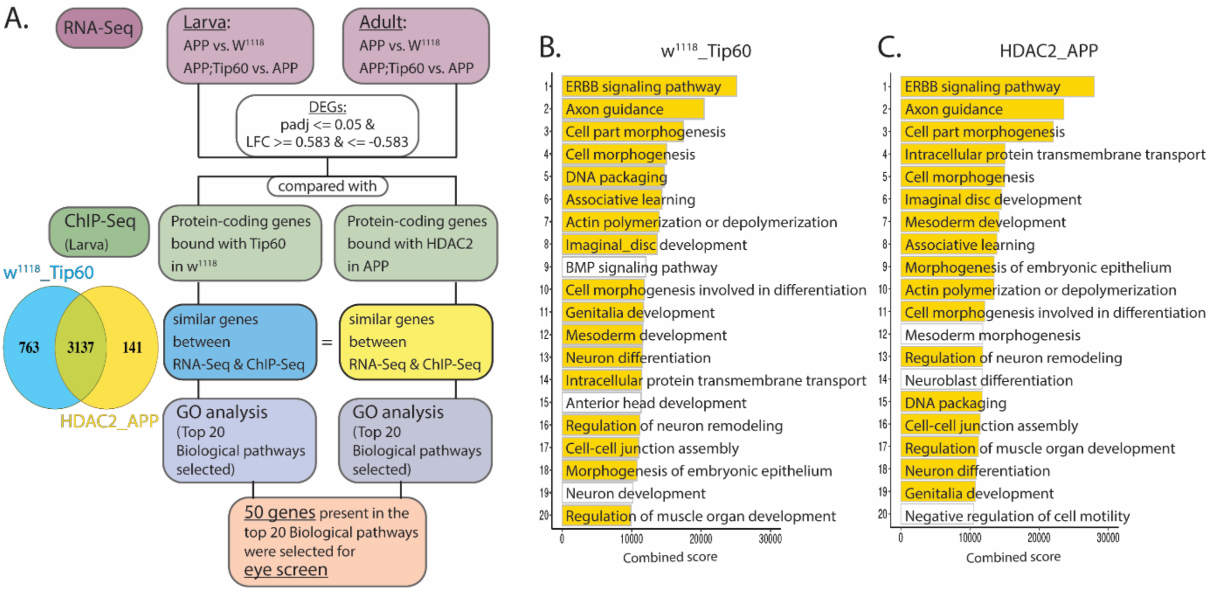


**Figure S5:** (A) Flow chart representing the gene selection process for the eye screen. The differentially regulated genes (RNA-seq) were compared with the protein-coding genes bound by Tip60 in w1118 genotype and by HDAC2 in APP genotype. Venn diagram represents the overlap between genes bound by Tip60 in w1118 and by HDAC2 in APP. 50 genes, bound by both HDAC2 and Tip60, were randomly selected from the top 20 biological pathways enriched by the gene set of both Tip60 (B) and HDAC2(C) and Tip60.

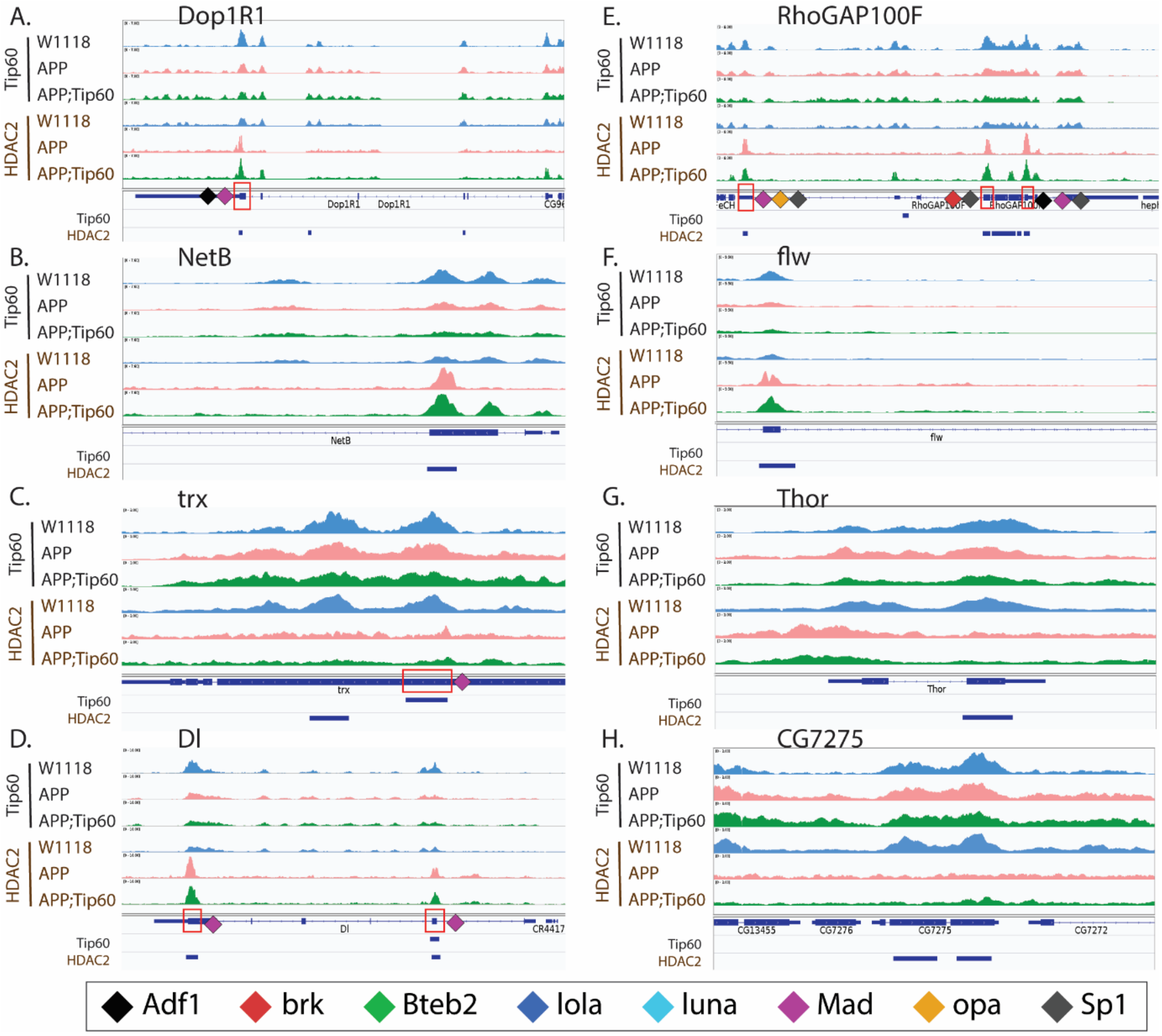


**Figure S6:** (A-H) Genome browser track view of Tip60 and HDAC2 peaks in three genotypes ($w^{1118}$, APP, and APP;Tip60) at the Dop1R1 (A), NetB (B), trx (C), Dl (D), RhoGAP100F (E), flw (F), Thor (G), and CG7275 (H) loci. Below the tracks, the gene features panel has loci marked: representing the transcription factor (Adf1, brk, Bteb2, lola, luna, Mad, opa, and Sp1) binding sites. The blue bars below the gene features panel depicts the regions bound by Tip60 and HDAC2. These genes with significantly enriched peaks exhibit a prominent phenotypical difference in eye screen.

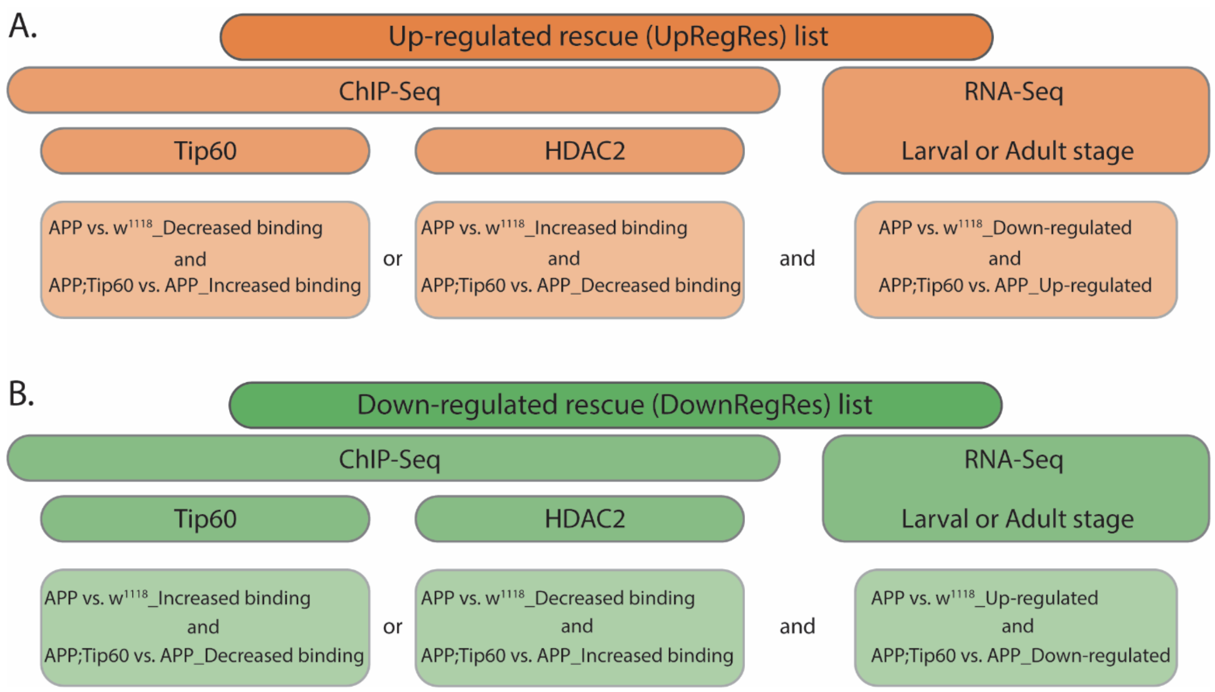


**Figure S7: Schematic representation of the selection process to obtain gene list considered as rescue list (Up-regulated and down-regulated rescue list).** The selection procedure is based on the concept that an increase in Tip60 binding or decrease in HDAC2 binding on genes (ChIP-seq) results in their increased expression (RNA-seq) and vice versa. The gene list of APP-induced binding of HDAC2 or Tip60 reversed by the Tip60 expression (ChIP-seq) is compared with the gene list of APP-induced genes reversed by Tip60 expression in either larval or adult heads (RNA-seq). The genes present in both the list (ChIP-seq and RNA-seq) are considered as rescue list.

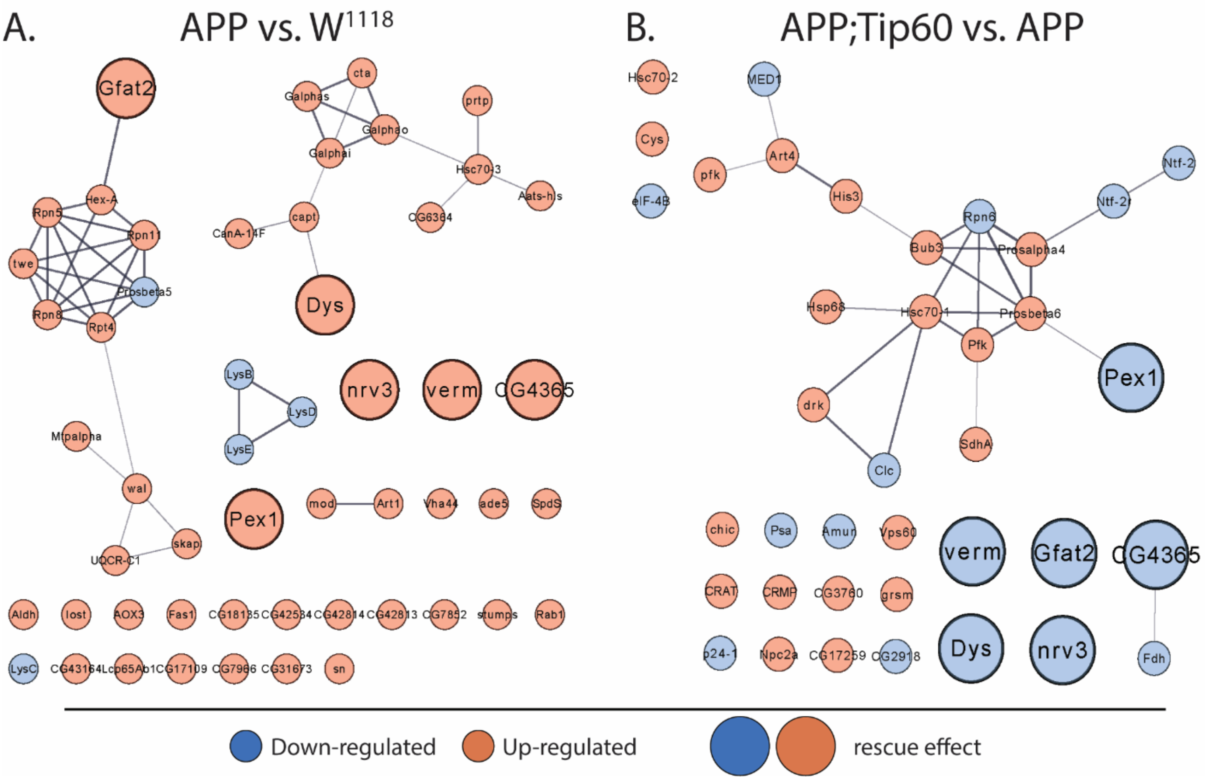


**Figure S8: A STRING network analysis of the proteins regulated by the APP and Tip60 expression in the third instar larval heads.** Protein networks of (A) APP-induced (APP vs. $w^{1118}$) and (B) Tip60 regulated (APP;Tip60 vs. APP) proteins. Networks of all differentially regulated proteins with a STRING interaction medium confidence (0.4). Here, network nodes represent proteins, and edges represent protein-protein associations. Blue-colored circles represent down-regulation, and red-colored circles represent up-regulation. Large circles represent APP-induced proteins reversed by the Tip60 expression (rescue effect).
